# Acute COG inactivation unveiled its immediate impact on Golgi and illuminated the nature of intra-Golgi recycling vesicles

**DOI:** 10.1101/2022.05.24.493317

**Authors:** Farhana Taher Sumya, Irina D. Pokrovskaya, Zinia D’Souza, Vladimir V. Lupashin

## Abstract

Conserved Oligomeric Golgi (COG) complex controls Golgi trafficking and glycosylation, but the precise COG mechanism is unknown. The auxin-inducible acute degradation system was employed to investigate initial defects resulting from COG dysfunction. We found that acute COG inactivation caused a massive accumulation of COG-dependent (CCD) vesicles that carry the bulk of Golgi enzymes and resident proteins. v-SNAREs (GS15, GS28) and v-tethers (giantin, golgin84, and TMF1) were relocalized into CCD vesicles, while t-SNAREs (STX5, YKT6), t-tethers (GM130, p115), and most of Rab proteins remained Golgi-associated. Airyscan microscopy and velocity gradient analysis revealed that different Golgi residents are segregated into different populations of CCD vesicles. Acute COG depletion significantly affected three Golgi-based vesicular coats- COPI, AP1, and GGA, suggesting that COG uniquely orchestrates tethering of multiple types of intra-Golgi CCD vesicles produced by different coat machineries. This study provided the first detailed view of primary cellular defects associated with COG dysfunction in human cells.

## Introduction

Newly synthesized proteins are delivered from the endoplasmic reticulum (ER) to Golgi for processing, sorting, and secretion (Ungar *et al*., 2006; I. Pokrovskaya *et al*., 2011; Cottam and Ungar, 2012; Willett, Ungar and Lupashin, 2013; Huang and Wang, 2017; D’Souza, Taher and Lupashin, 2020; D’Souza *et al*., 2021). Glycosylation is one of the major cargo modifications chiefly carried out by the Golgi (Stanley, 2011). Modification of glycoconjugates requires the transfer of sugar donors onto acceptor substrates (proteins and lipids) by glycosyltransferases and partial remodeling by glycosidases. Under the cisternal maturation model, secretory and transmembrane cargo molecules remain in the lumen of the Golgi cisternae while the cisternae themselves progress through the stack and ‘mature’ through recycling of their resident proteins (Glick and Nakano, 2009). During maturation, each Golgi cisterna needs to maintain its specific set of glycosylation enzymes, sugar transporters, and other cargo modifiers. Proper cisternal compartmentalization of the glycosylation machinery is vital as glycosylation is template independent (Berninsone and Hirschberg, 2000; Stanley, 2011; D’Souza, Taher and Lupashin, 2020; D’Souza *et al*., 2021). There are continuous discussions on the exact mechanisms and pathways used by different cells and organisms for the anterograde cargo transport through the Golgi (Orci, Ravazzola, *et al*., 2000; Pelham, 2001; Hwang, 2008; Mironov and Beznoussenko, 2019), but the majority of recent studies agree on general rules for the maintenance of Golgi enzymes. It is reported that both active retention and recycling mechanisms are utilized to maintain the proper localization of Golgi resident proteins (Glick and Nakano, 2009; Rizzo *et al*., 2013). The recycling of Golgi resident proteins and enzymes is mostly facilitated by COPI vesicle-mediated retrograde transport (Pelham, 2001; Cottam and Ungar, 2012; Di Martino, Sticco and Luini, 2019; D’Souza *et al*., 2021; Park *et al*., 2021). A balance between anterograde and retrograde transport is not only important for maintaining proper concentration of the resident Golgi proteins and lipids but also for the cell’s physiology (Blackburn, D’Souza and Lupashin, 2019; D’Souza, Taher and Lupashin, 2020). The vesicular trafficking machinery consists of several distinct modules that drive vesicle budding from a donor compartment followed by its transport, tethering, and fusion with the acceptor compartment (Cai, Reinisch and Ferro-Novick, 2007; Cottam and Ungar, 2012; D’Souza *et al*., 2021). Vesicle formation is initiated by small ARF family GTPases that recruit coat proteins which collect and segregate cargo molecules into 60 mm vesicles. The vesicle then buds off, gets uncoated, and then specifically tethered to the target membrane. Vesicle tethering is achieved by both coiled-coil and MTC (multisubunit tethering complex) tethers. followed by vesicle fusion with a specific Golgi subcompartment in a SNARE-dependent reaction (Willett, Ungar and Lupashin, 2013; Blackburn, D’Souza and Lupashin, 2019).

The conserved oligomeric Golgi (COG) complex is the major Golgi MTC (Willett, Ungar and Lupashin, 2013; Blackburn, D’Souza and Lupashin, 2019; D’Souza *et al*., 2019; D’Souza, Taher and Lupashin, 2020; D’Souza *et al*., 2021). COG is composed of 8 subunits COG1-COG8 (Ungar *et al*., 2002; Lees *et al*., 2010; Blackburn, D’Souza and Lupashin, 2019; D’Souza, Taher and Lupashin, 2020; D’Souza *et al*., 2021), which are organized in two subcomplexes, lobe A and lobe B (Fotso *et al*., 2005, ; Ungar *et al*., 2005). COG orchestrates retrograde intra-Golgi vesicular trafficking by tethering vesicles carrying recycling Golgi resident proteins (such as glycosylation enzymes and nucleotide sugar transporters) back to their working compartments thereby facilitating proper glycosylation of secretory and transmembrane proteins (Shestakova, Zolov and Lupashin, 2006; Steet and Kornfeld, 2006; I. Pokrovskaya *et al*., 2011; Willett *et al*., 2014;). To achieve its role in membrane trafficking, COG physically and functionally interacts with other components of vesicular trafficking machinery, including SNAREs, SNARE-interacting proteins, Rabs, coiled-coil tethers, and coat proteins (Ungar *et al*., 2006; Willett, Ungar and Lupashin, 2013; Blackburn, D’Souza and Lupashin, 2019; D’Souza *et al*., 2021). COG malfunction in humans causes global glycosylation defects termed COG-related Congenital Disorders of Glycosylation (COG-CDGs) (Foulquier, 2009; D’Souza, Taher and Lupashin, 2020; Sumya, Pokrovskaya and Lupashin, 2021). COG-CDGs are multi-systemic disorders with several common symptoms including global developmental defects, dysmorphic features, microcephaly, and failure to thrive which are accompanied by the liver and neurological impairment. It has been reported that almost one-third of the patients with congenital defects in the Golgi glycosylation have mutations in COG complex subunits. More than 30 different COG mutations have been identified to date (Zeevaert *et al*., 2008; Foulquier, 2009; Reynders *et al*., 2011; D’Souza, Taher and Lupashin, 2020; D’Souza *et al*., 2021; Ondruskova *et al*., 2021).

Multiple knockouts (KO), knock-down (KD), and knock-sideways approaches have been applied to unravel the details of the COG’s cellular functions (Kingsley *et al*., 1986; Zolov and Lupashin, 2005; I. Pokrovskaya *et al*., 2011; Laufman, Hong and Lev, 2013; Willett *et al*., 2013, 2016; Bailey Blackburn *et al*., 2016; Blackburn and Lupashin, 2016; Climer *et al*., 2018; Petitjean *et al*., 2020). The complete KO of individual COG subunits in HEK293T cells resulted in abnormal Golgi morphology, accumulation of Enlarged Endo-Lysosomal Structures (EELSs), inhibited retrograde protein trafficking, and altered the repertoire of secreted proteins (Blackburn *et al*., 2018; D’Souza *et al*., 2019). At the same time, the COG KD study revealed a massive accumulation of COG complex dependent (CCD) vesicles that carry Golgi enzymes and intra-Golgi v-SNAREs (Zolov and Lupashin, 2005; Shestakova, Zolov and Lupashin, 2006). Both COG mutations, KO or KD of COG subunits resulted in altered glycosylation of both N-, and O-linked glycans as well as destabilization or mislocalization of Golgi glycosylation machinery (Ungar *et al*., 2002; Oka *et al*., 2005; Zolov and Lupashin, 2005; I. Pokrovskaya *et al*., 2011; D’Souza, Taher and Lupashin, 2020). Though CRISPR-based KO and RNAi-interference are important and robust approaches to studying COG function, they both require a relatively long time (3-10 days) to produce a mutant phenotype; KO and KD experiments can result in incomplete silencing, off-target effects and adaptation, therefore observed mutant phenotypes could be either direct or indirect consequences of COG depletion. Golgi membranes are highly dynamic and transport through the Golgi in human cells usually takes less than 30 minutes (Boncompain *et al*., 2012; Beznoussenko *et al*., 2014), so we reasoned that rapid silencing of COG subunits would bring us closer to precisely elucidating COG’s role in intra-Golgi trafficking..

In this study, we have created a novel cellular system to investigate the immediate effect of rapid COG depletion on Golgi physiology. Auxin inducible degron (AID) approach (Nishimura *et al*., 2009; Holland *et al*., 2012; Zhang and Seemann, 2021) has been applied to completely degrade the COG4 subunit in RPE1 cells within 30 min after adding auxin. This 30 min period was comparable with the trafficking time through the Golgi allowing visualization of instant effects of COG dysfunction on Golgi structure and dynamics. We applied a combination of biochemical and microscopic approaches to dissect the impact of the acute COG4 depletion on other COG subunits, COG interacting membrane trafficking partners, glycosylation enzymes, as well as other Golgi resident proteins. This study provided the first detailed view of primary cellular defects associated with COG dysfunction in human cells.

## Materials and Methods

### Cell Culture and Auxin treatment

hTERT RPE1 (Retinal Pigment Epithelial) and HEK293T cells were purchased from ATCC. hTERT RPE1 COG4 KO cells were described previously (PMID: 34603392). Cells were cultured in Dulbecco’s Modified Eagle’s Medium (DMEM) containing Nutrient mixture F-12 (DMEM/F12, Corning 10–092-CV) supplemented with 10% Fetal Bovine Serum (Atlas Biologicals, CF-0500-A). Cells were incubated in a 37°C incubator with 5% CO_2_ and 90% humidity.

For rapid COG4 degradation, a stock solution of 0.5 M Indole-3-acetic acid sodium salt (auxin, IAA, Sigma # I5148) was prepared in water and stored in a frozen aliquot. Time course treatment of cells was performed with 500 µM IAA for 0.5, 1, 2, 24, and 48 hours at 37°C. The cells without auxin treatment were considered as untreated control.

### Plasmid preparation, generation of lentiviral particles, and stable cell lines

All constructs were developed using standard molecular biology techniques and are listed in **Table 1**

| Plasmid Name | Source | Citation |
| --- | --- | --- |
| pBabe OsTIR1-9myc (PuroR) | Addgene #47328 | (Holland *et al.*, 2012) |
| pMK292 mAID-mCherry2-NeoR | Addgene #72830 | (Natsume *et al.*, 2016) |
| pUMVC (packaging plasmid for producing MNuLV retroviral particles) | Addgene #8449 | (Stewart *et al.*, 2003) |
| pMD2.G (VSV-G envelope expressing plasmid) (a gift from Didier Trono) | (Addgene plasmid # 12259; http://n2t.net/addgene:12259 ; RRID: Addgene_12259) | (Dull *et al.*, 1998) |
| pRSV-Rev | Addgene #12253 | (Dull *et al.*, 1998) |
| pMDLg/pRRE | Addgene #12251 | (Dull *et al.*, 1998) |
| COG4-2xGFP in pEntra1A | Lab | (Sumya, Pokrovskaya and Lupashin, 2021) |
| pLenti COG4 Neo DEST plasmid with COG4 promoter | Lab | (Khakurel *et al.*, 2021; Sumya, Pokrovskaya and Lupashin, 2021) |
| COG4-mAID-mCherry in pENTRA1A | This study | - |
| COG4-mAID-mCherry in pLenti | This study | - |

### Generation of Retroviral Particles and COG4 KO Cell line Expressing OsTIR1

OsTIR1-9myc was stably expressed in COG4KO cells to induce depletion of AID-tagged COG4 protein. pUMVC (5.2 µg), pMD2.G (2.6 µg), and pBabe OsTIR1-9myc (7.2 µg) were mixed to transfect HEK293FT cells using Lipofectamine 3000 using a standard protocol. Transfected HEK293FT cells were placed in serum-reduced Opti-MEM with 25 μM Chloroquine and 1x GlutaMAX. 5 hours after transfection Na-Butyrate (5 mM final concentration) was added. The next day, the medium was changed to Opti-MEM supplemented with 1x GlutaMAX. At 48 h after transfection, the medium was collected, and cell debris was removed by centrifugation at 1000×g for 5 min. The supernatant was clarified by passing through a 0.45 μM polyethersulfone (PES) membrane filter; the viral supernatant was frozen in 1 ml aliquots and stored at −80°C.

hTERT RPE1 COG4 KO cells were plated on a 6-wells plate in DMEM/F12 at 50% confluency. The next day, cells were transduced with 500 µl of viral supernatant. 48 h after transduction the retroviral media was exchanged for fresh DMEM/F12 growth media containing Puromycin (10 µg/ml final concentration, selection dose). After 48 h of puromycin selection, the media was replaced with complete media containing 5 µg/ml of puromycin (maintenance dose). The single-cell clones were isolated into 96 well plates by serial dilution. Cells were allowed to grow for two more weeks before expanding. After that, cell colonies were collected by trypsin detachment and expanded into a 12-well plate with a complete media containing the maintenance dose of puromycin. Expanded clones were screened by western blot (WB) and immunofluorescent (IF) analysis to identify the cells with uniform OsTIR1-9myc expression.

### Construction of Plasmid COG4-mAID-mCherry in pENTRA1A

To produce COG4-mAID-mCherry in pENTRA1A, mAID-mCherry was first amplified by PCR from pMK292 mAID-mCherry2-NeoR plasmid (Addgene #72830) using primers 5’- AATTGGTACCGGATCCGGTGCAGGCGCCAAG-3’, and 5’- GCGCCTCGAGTTACTTGTACAGCTCGTCGTCCAT-3’ following KpnI/XhoI digestion and ligation of PCR fragment with similarly digested COG4-2xGFP in pEntra1A (Sumya, Pokrovskaya and Lupashin, 2021).

### Production of COG4-mAID-mCherry lentivirus and COG4 KO-OsTIR1 expressing AID tagged COG4 stable cell line

COG4-mAID-mCherry in pEntra1A were recombined into pLenti COG4 Neo DEST plasmid with COG4 promoter (Khakurel *et al*., 2021; Sumya, Pokrovskaya and Lupashin, 2021) to generate COG4-mAID-mCherry in pLenti using Gateway LR Clonase II Enzyme Mix (Thermo Fisher) and transformed into Stbl3 competent cells according to the manufacturer’s instructions. DNA was extracted with QIAprep Spin Miniprep DNA extraction Kit. Correct COG4-mAID-mCherry pLenti clones were tested by restriction analysis. COG4-mAID-mCherry expression was validated by transfecting HEK293T cells with selected COG4-mAID-mCherry pLenti plasmids followed by WB analysis of total cell lysates using COG4 antibody. To produce lentiviral particles, equal amounts of lentiviral packaging plasmids pMD2.G [a gift from Didier Trono (Addgene plasmid #12259; http://n2t.net/addgene:12259; RRID: Addgene_12259)], pRSV-Rev, pMDLg/pRRE (Dull *et al*., 1998), and COG4-mAID-mCherry pLenti were mixed to transfect HEK293FT cells with Lipofectamine 3000 using a manufacturer protocol. Transfected HEK293FT cells were placed in serum-reduced Opti-MEM supplemented with 25 μM Chloroquine and 1x GlutaMAX. The next day, the medium was changed to Opti-MEM supplemented with 1x GlutaMAX. At 72 h after transfection, the medium was collected, and cell debris was removed by centrifugation at 600×g for 10 min. The supernatant was passed through a 0.45 μM polyethersulfone (PES) membrane filter and the lentiviral medium was stored at 4°C overnight or splitted into aliquots, snap freezed in liquid nitrogen, and stored at −80°C.

hTERT RPE1 COG4 KO OsTIR1-9myc cells were plated in two wells of a 6-wells plate in complete medium to reach 90% confluency the next day. One of the wells was used as a control for antibiotic selection. The next day, cells were transduced with 500 µl of lentiviral supernatant. At 48 h after transduction, the lentiviral media was substituted to cell growth media containing G418 (600 µg/ml final concentration, selection dose). After 48 h of selection, the media was replaced with complete media containing 200 µg/ml of G418 (maintenance dose). The cells were expanded at 37°C and 5% CO_2_ for 48 h. After G418 selection, the single-cell clones were isolated into 96 well plate by serial dilution. Cells were allowed to grow for two weeks, collected by trypsin treatment, and expanded each colony into a 12-well plate with a complete medium containing G418. WB and IF analyses were performed to identify the clone with COG4-mAID-mCherry expression. Clones with a uniformed expression of COG4-mAID-mCherry were split onto 10-cm dishes, aliquots were cryopreserved in 2x freezing medium (80% FBS with 20% DMSO) mixed with growth medium.

### Preparation of Cell Lysates and Western Blot Analysis

To prepare the cell lysates, cells grown on tissue culture dishes were washed three times with phosphate-buffered saline (PBS) and lysed in hot 2% SDS. Samples were mixed with 6x SDS sample buffer containing β-mercaptoethanol and heated for 10 min at 70°C. To prepare the lysates for each fraction mentioned in the membrane fractionation experiment, membrane pellets were resuspended in 2% SDS following the addition of 6xSDS sample buffer containing β-mercaptoethanol. For making the sample for supernatant (mentioned in the membrane fractionation experiment) and fractions of vesicle gradients 6xSDS sample buffer containing β-mercaptoethanol was added. The samples were heated for 5 min at 95°C following 10 min at 70°C.

10–20 µg of protein was loaded into Bio-Rad (4–15%) or Genescript (8–16%) gradient gel. Proteins were transferred onto nitrocellulose membrane using the Thermo Scientific Pierce G2 Fast Blotter. Membranes were washed in PBS, blocked in Bio-Rad blocking buffer for 20 min, and incubated with primary antibodies for 1 h at room temperature or overnight at 4°C. Membranes were washed with PBS and incubated with secondary fluorescently-tagged antibodies diluted in Bio-Rad blocking buffer for 1 h. All the primary and secondary antibodies used in the study are listed in **Table 2**. Blots were then washed with PBS and imaged using the Odyssey Imaging System. Images were processed using the LI-COR Image Studio software.

**Table 2.**
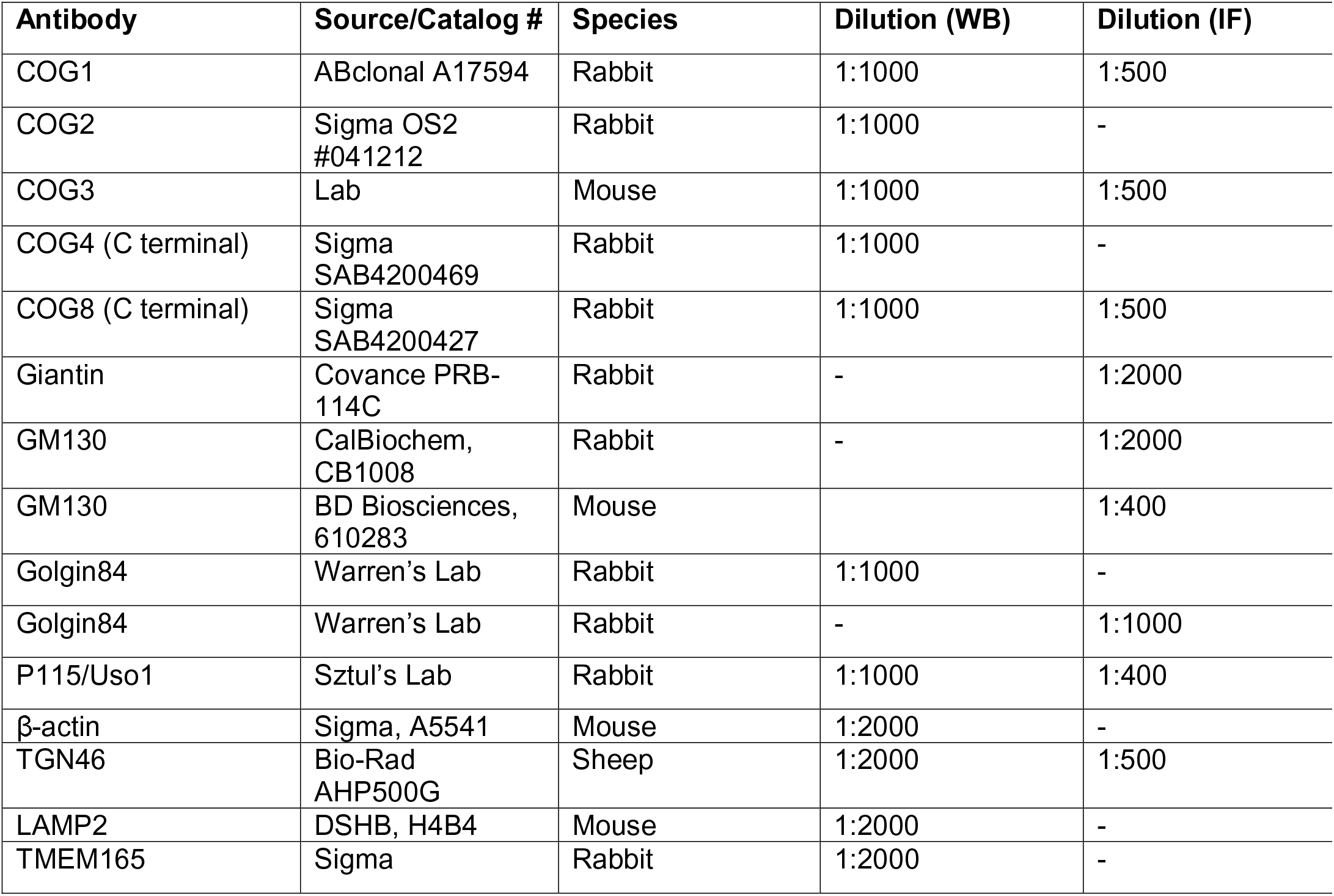
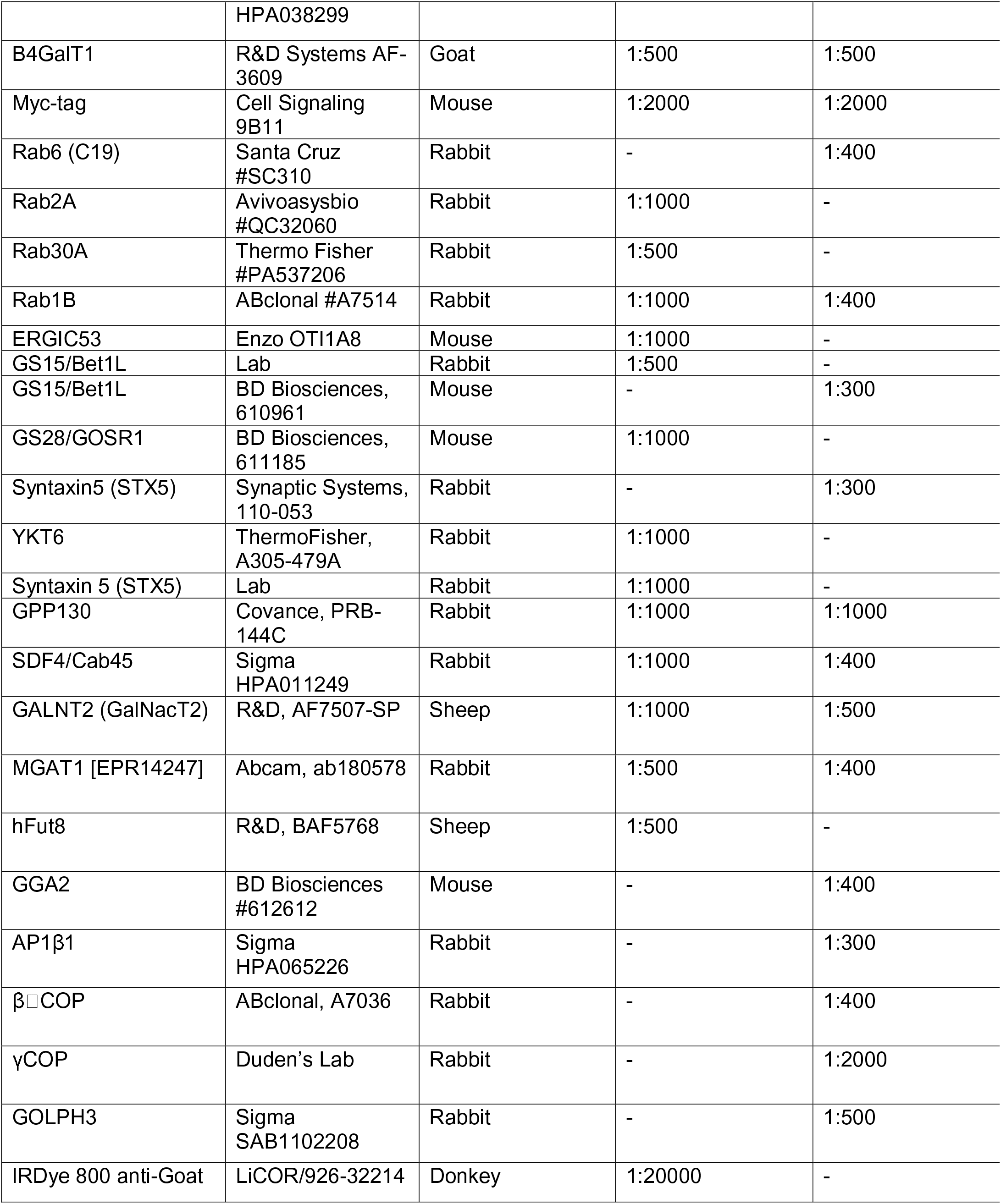
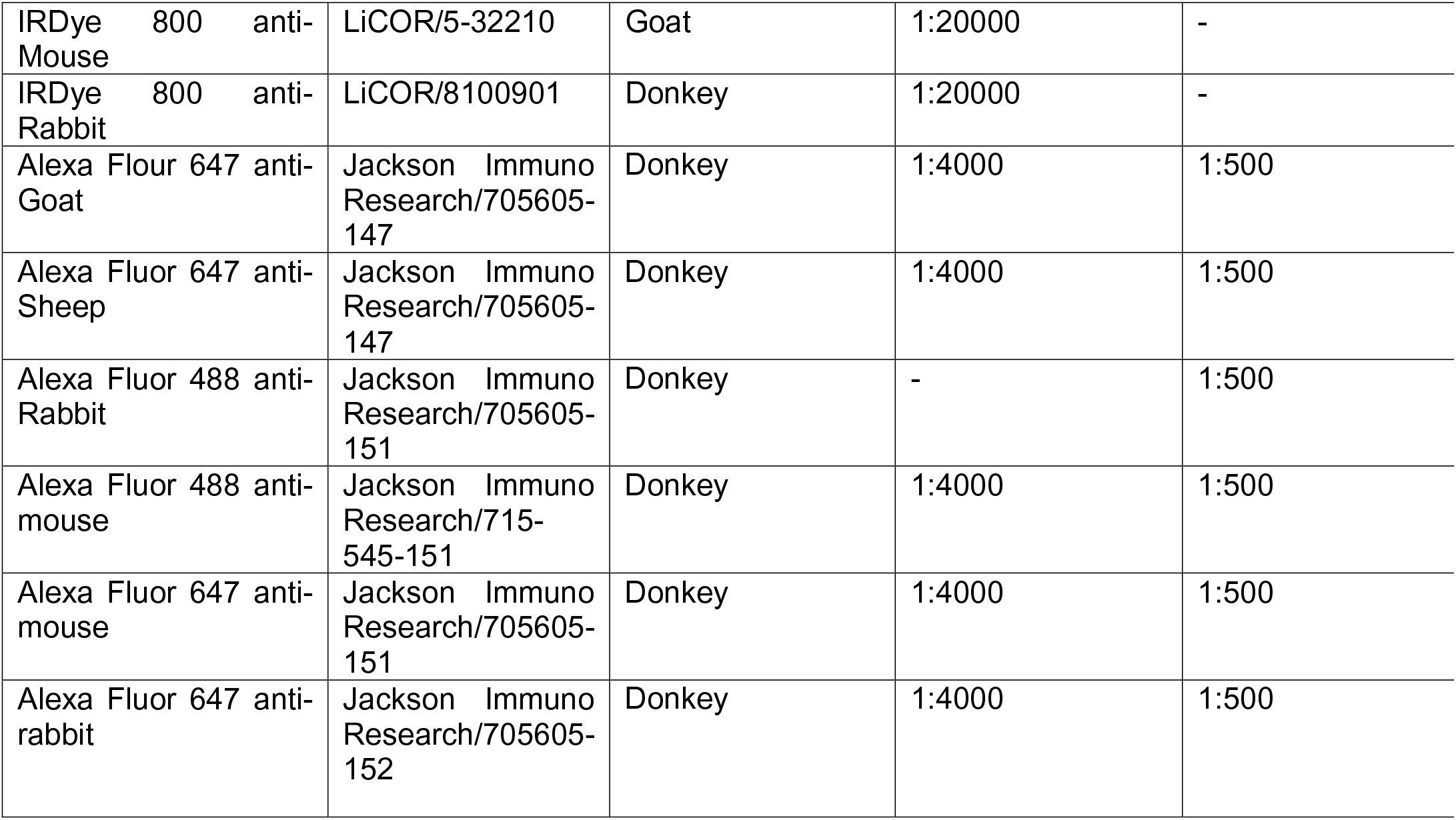
List of antibodies

### Lectin Blotting

To perform blots with fluorescent lectins, 10 µg of cell lysates were loaded onto Bio-Rad (4– 15%) gradient gels, and proteins were transferred to nitrocellulose membrane using the Thermo Scientific Pierce G2 Fast Blotter. The membranes were blocked with Bio-Rad blocking buffer for 20 min. Helix Pomatia Agglutinin (HPA)-Alexa 647 (Thermo Fisher) or Galanthus Nivalis Lectin (GNL) conjugated to Alexa 647 (Blackburn and Lupashin, 2016) were diluted 1:1,000 in Bio-Rad blocking buffer from their stock concentration of 1 µg/µl and 5 µg/µl, respectively. Membranes were incubated with diluted HPA-647 or GNL-647 for 1 h, washed four times in PBS and imaged using the Odyssey Imaging System.

### Superresolution AiryScan Fluorescent Microscopy

Cells were grown on 12-mm round coverslips to 80–90% confluency were fixed with 4% paraformaldehyde (PFA, freshly made from 16% stock solution) diluted in PBS for 15 min at room temperature. Cells were then permeabilized with 0.1% Triton X-100 for 1 min followed by treatment with 50 mM ammonium chloride for 5 min, treated with 6 M urea for 2 min (only for COG3 staining), and washed twice with PBS. Blocking (two incubations 10 min each) in 1% BSA, 0.1% saponin in PBS was done. Cells were then incubated with primary antibody (diluted in 1% cold fish gelatin, 0.1% saponin in PBS) for 45 min, washed, and incubated with fluorescently conjugated secondary antibodies diluted in the antibody buffer for 30 min. Cells were washed four times with PBS, then coverslips were dipped in PBS and water 10 times each and mounted on glass microscope slides using Prolong^®^ Gold antifade reagent (Life Technologies). Cells were imaged with a 63× oil 1.4 numerical aperture (NA) objective of an LSM880 Zeiss Laser inverted microscope with Airyscan using ZEN software.

### Analysis of Golgi fragmentation

Ten fields (at least 30 individual cells) of GM130-stained Airyscan microscopic images of untreated (control) or auxin treated (IAA) COG4-mAID cells were used. ImageJ software was used to create binaries. Then the Golgi particles having a surface area <1µm^2 were counted using the ‘Analyze Particle’ function of ImageJ. These particles were considered “Golgi fragments”. The average number of fragmented Golgi in auxin-treated COG4-mAID cells was compared with untreated control.

### Membrane Fractionation Experiment

Cells grown in 15 cm dishes to 90% confluency, were washed with PBS and collected by trypsinization followed by centrifugation at 400xg for five minutes. The cells pellet was resuspended in 1.5 ml of cell collection solution (0.25 M sucrose in PBS) followed by centrifugation at 400xg for five minutes. The pellet was then resuspended in 1.5 ml of a hypotonic lysis solution (20 mM HEPES pH 7.2, with protein inhibitor cocktail, and 1 mM PMSF) and passed through a 25 G needle 20 times to disrupt cells. Cell lysis efficiency was evaluated under the phase-contrast microscope. After that KCL (to 150 mM final concentration) and EDTA (2 mM final) were added. Unlysed cells and cell nuclei were removed by centrifugation at 1000xg. The post-nuclear supernatant (PNS) was transferred to the 1.5 ml Beckman tube (#357488) and the Golgi-enriched fraction was sedimented at 30,000xg for 10 minutes. The Supernatant (S30) was transferred into a new Beckman tube and the vesicle-enriched fraction was pelleted at 100,000xg for 1 hour, at 4°C using a TLA-55 rotor. The samples from each fraction were prepared to perform WB analysis.

### Fractionation of Vesicles by Velocity Sedimentation

Fractionation of vesicles by velocity sedimentation was done following a published protocol (Love *et al*., 1998) with some modifications. Before placing the sucrose fraction Beckman ultra-clear 2 mL centrifuge tube (347356) was coated with siliconizing reagent Sigmacote (Sigma Aldrich), rinsed with water, and dried for 1 h. 200 µl each of 35, 32.5, 30, 27.5, 25, 22.5, 20, 17.5, 15, and 12.5% (wt/wt) sucrose in KHM buffer (150 mM KCl, 10 mM Hepes-KOH, pH 7.2, 2.5 mM MgOAc) were overlaid in a siliconized centrifuge tube and left at room temperature for two hours to create a linear gradient. Vesicular pellet (P100) was resuspended in KHM buffer. 200 µl of resuspended vesicle fraction was laid on top of sucrose gradients and the gradient was centrifuged for 1 h at 186,000xg in an ultracentrifuge rotor TLS55 at 4°C. Eleven fractions of 200 µl each were manually collected from the top. The samples were mixed with 6x sample buffer and prepared for WB analysis.

### High-pressure freezing, freeze substitution, and EM

#### High-pressure freezing (HPF)/FS (Freezing substitution)

Sapphire disks were coated with a 10 nm carbon layer followed by collagen (Corning) coating per the manufacturer’s instructions. Coated disks were sterilized under UV light and transferred to new sterile 3 cm dishes for plating the cells. After reaching 80– 100% confluence, cells were equilibrated in fresh media for 2–3 h at 37°C, treated with Auxin various times. High-pressure freezing at specified time points was performed in cryo-protectant (PBS with 2% Type IX ultra-low melt agarose (Sigma-Aldrich), 100 mM D-mannitol, and 2% FBS) using a Leica EM PACT2 high-pressure freezing unit (Leica Microsystems) with rapid transfer system at high pressure (2100 bar). All solutions, bayonets, and sample holders were pre-warmed to 37°C, all manipulations were carried out on a 37°C heating platform.

#### Freeze substitution dehydration

Samples were transferred under liquid nitrogen to cryovials containing anhydrous acetone with 2% osmium tetroxide, 0.1% glutaraldehyde, and 1% double-distilled (dd) H_2_O. Next, the tubes were transferred to a freeze-substitution chamber at –90°C programmed with the following schedule: –90°C for 22 h, warm 3°C/h to –60°C, –60°C for 8 h, warm 3°C/h to –30°C, –30°C for 8 h, warm 3°C/h to 0°C. Afterward, sample tubes were placed on ice and moved to the cold room (4°C). After washing three times with acetone samples were stained with a 1% tannic acid, 1% ddH_2_O solution in acetone on ice for 1 h followed by three acetone washes. Next, samples were stained with a 1% OsO_4,_ 1% ddH_2_O solution in acetone on ice for 1 h, washed 3 x 10 min in acetone, and dehydrated over a series of ethanol gradations (25%, 50%, 75%, 100%) using automatic resin infiltration protocol for PELCO Bio-Wave Pro laboratory microwave system. Samples were embedded in Araldite 502/Embed 812 resins with a DMP-30 activator and baked at 60°C for 48 h.

#### Thin section TEM

Thin sections were cut at a thickness of 50 nm with a Leica UltraCut-UCT microtome and post-stained with aqueous uranyl acetate and Reynold’s lead citrate (EMS). <u>Electron microscopy and image handling:</u> Images were taken using an FEI Tecnai TF20 intermediate-voltage electron microscope operated at 80 keV (FEI Co.). Images were acquired with an FEI Eagle 4k digital camera controlled with FEI software.

### Antibodies

Primary and secondary antibodies used for WB or IF were made in the lab, received from colleagues or commercially purchased. The list of antibodies and their dilutions are described in **Table 2**.

### Statistical Analysis

All the WB images are representative of 3 repeats and those were quantified by densitometry using the LI-COR Image Studio software. Error bars for all graphs represent standard deviation. Statistical analysis was done using one-way ANOVA or paired *t-*test using GraphPad Prism software. In the case of IF analysis, each dot in the bar graph represents the colocalization of GM130 and other Golgi proteins in several (1 to 10) cells imaged per field.

## Results

### Establishment of cellular system for the acute depletion of COG4

To achieve rapid inactivation of the COG complex we employed mAID degron tagging of the COG4 subunit in combination with co-expression of auxin perceptive F-box protein OsTIR1 (Dharmasiri and Estelle, 2004; Nishimura *et al*., 2009; Zhang and Seemann, 2021). COG4 is an essential subunit of lobe A COG subcomplex that interacts with key elements of the vesicle fusion machinery: STX5, Sly1/SCFD1, and Rab30 (Kim *et al*., 2001; Shestakova *et al*., 2007; Laufman *et al*., 2009; Laufman, Hong and Lev, 2013; Miller *et al*., 2013; Willett *et al*., 2013, 2014; Willett, Ungar and Lupashin, 2013) and we reasoned that COG4 inactivation will be sufficient to compromise COG complex functions. To induce auxin-mediated COG4 degradation, RPE1 COG4 knockout (KO) cells (Sumya, Pokrovskaya and Lupashin, 2021) were sequentially transduced with the retroviral construct expressing OsTIR1-9myc and the lentiviral construct expressing COG4-mAID-mCherry (COG4-mAID) hybrid protein under the control of endogenous COG4 promoter **(Figure 1A)**. After the addition of the auxin (Indole-3-acetic acid/IAA), the COG4-mAID protein should become polyubiquitinated and degraded by the proteasome **(Figure 1A)**. The resulting cell line RPE1-COG4-mAID was viable, exhibiting growth and morphological characteristics similar to the wild-type RPE1 cells (FTS, VL, unpublished observation). Next, the expression and functionality of COG4-mAID were tested by using WB and IF approaches. The results revealed that COG4-mAID was expressed at a near endogenous level compared to COG4 expression in wild-type cells **(Figure 1B)**. COG4-mAID protein was Golgi-localized in the perinuclear region and colocalized with Golgi marker GM130/GOLGA2 **(Figure 1C, upper panel)**. WB analysis of COG4-mAID cells validated the expression of TIR-9myc essential for auxin-induced degradation of COG4-mAID **(Figure 1B)**. Previously we have reported that the stability and glycosylation of trans-Golgi enzyme B4GalT1, putative Mn2+ transporter TMEM165, cis-Golgi recycling glycoprotein GPP130/GOLIM4 and lysosomal glycoprotein Lamp2 are altered in COG KO HEK293T cells (Blackburn *et al*., 2018; Sumya, Pokrovskaya and Lupashin, 2021). WB analysis of RPE1 COG4 KO cells confirms depletion of B4GalT1 and GPP130, as well as an increase in the electrophoretic mobility of TMEM165 and LAMP2, indicating defects in glycosylation **(Figure 1B)**. The expression of COG4-mAID rescued the stability and glycosylation of all affected proteins **(Figure 1B),** suggesting that the COG4-mAID is functional and rescued the major biochemical phenotypes associated with COG deficiency. **(Figure 1B)**.

**Figure 1.**
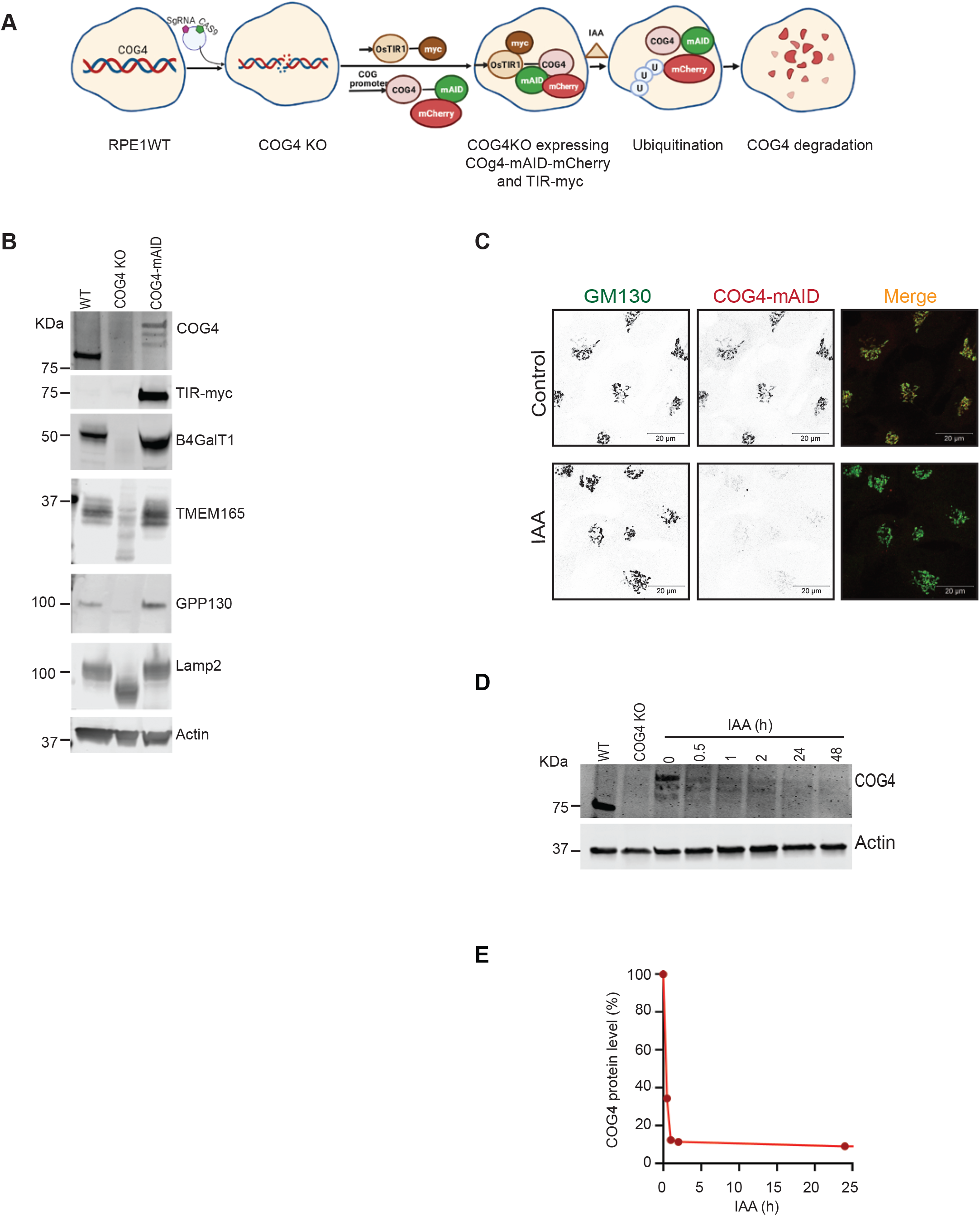
COG4-mAID-mCherry (COG4-mAID) is functionally substituted endogenous COG4 and rapidly depleted upon IAA (auxin) treatment. **(A)** The diagram shows the development of the RPE1 COG4 KO cell line co-expressing COG4-mAID-mCherry (COG4-mAID) under COG4 promoter and OsTIR1-9myc. The COG4-mAid is ubiquitinated and degraded upon IAA treatment. **(B)** Expression of COG4-mAID rescues major cellular phenotypes associated with COG4 deficiency. WB shows the expression of COG4, myc, and COG sensitive proteins in wild type, COG4 KO, and COG4-mAID cell lines. 10 µg of total cell lysates were loaded for each line. β actin has been used as a loading control. **(C)** COG4-mAID is Golgi localized (upper panel) and it is absent from the Golgi upon one-hour treatment with IAA. Airyscan superresolution IF analysis of COG4-mAID-mCherry (red) cells stained for GM130 (green). For better presentation green and red channels are shown in inverted black and white mode whereas the merged view is shown in RGB mode. Scale bars, 20 µm. **(D)** WB of time-dependent depletion of COG4-mAID upon IAA treatment. 10 µg of total cell lysates were loaded to each lane and probed with COG4 and actin antibodies. **(E)** The graph represents the quantification of D.

To test the IAA-induced degradation of COG4, the COG4-mAID cells were treated with IAA at five-time point intervals (0.5, 1, 2, 24, and 48 hours) and tested for COG4-mAID protein expression by WB. The result shows that IAA treatment leads to a significant decrease in the COG4-mAID protein level at 30 minutes and nearly complete COG4-mAID depletion after one hour of incubation with IAA **(Figure 1D)**. Airyscan superresolution microscopy revealed the absence of mCherry signal within one hour of IAA treatment **(Figure 1C lower panel).** Acute COG4 depletion did not significantly change the morphology of GM130-labeled Golgi and did not cause Golgi fragmentation, judged by the similar number of GM130 labeled Golgi fragments in control and IAA treated COG4-mAID cells **(Supplementary 1)**. As GM130 is not sensitive to COG4 depletion, it was used as a Golgi marker to check the colocalization of other Golgi proteins by IF.

### COG4 acute depletion affects the entire COG complex

To investigate the impact of acute COG4 depletion on other subunits of the COG complex, WB and IF approaches were applied. Airyscan microscopy revealed that COG1, COG3, and COG8 subunits were mislocalized from Golgi to the cytoplasm **(Figure 2A, B, C)** within one hour of COG4 depletion. As a result, the colocalization of COG subunits with GM130 was significantly decreased **(Figure 2D)**. Since the proteasomal degradation of COG4 resulted in off-Golgi localization of other COG subunits, we wonder if this would compromise their stability as well. Previously we showed that COG4 KO affects the total cellular level of COG2 and COG3 but has a lesser impact on lobe B subunits in the HEK293T cells (Bailey Blackburn *et al*., 2016). An efficient knockdown of COG3 also resulted in a reduction in COG1, 2, and 4 protein levels in HeLa cells (Zolov and Lupashin, 2005). We found that the acute depletion of COG4-mAID in RPE1 cells significantly affected the expression of COG1, COG2, and COG8, while the level of COG3 was affected only after a prolonged (48 h) depletion of COG4 **(Supplementary Figure 2)**. Together, the results suggest that acute COG4 depletion displaces the COG complex from the Golgi and reduces the cellular level of COG subunits.

**Figure 2.**
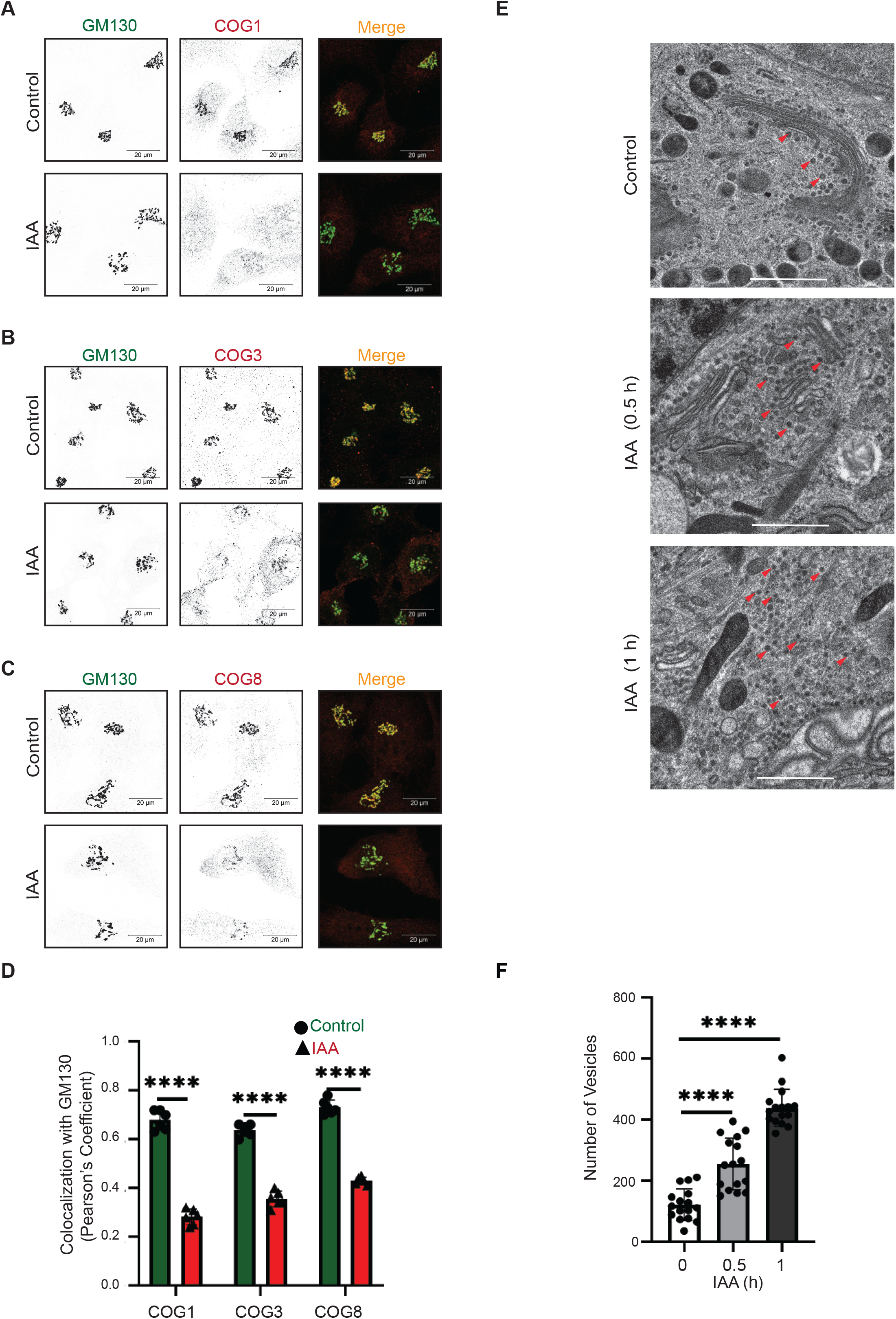
COG4 depletion affects other COG subunits and causes rapid accumulation of COG complex-dependent (CCD) vesicles. **(A)** Airyscan superresolution IF analysis of untreated (control) or IAA treated COG4-mAID cells stained for **(A)** GM130 (green) and COG1 (red), **(B)** GM130 (green) and COG3 (red) **(C)** GM130 (green) and COG8 (red). Scale bars, 20 µm. For better presentation green and red channels are shown in inverted black and white mode whereas the merged view is shown in RGB mode. **(D)** Colocalization of COG subunits with GM130 in control and IAA treated cells was determined using Pearson’s correlation coefficient and >60 cells were analyzed. Statistical significance was calculated by GraphPad Prism 8 using paired t-test. Here, ****P < 0.0001 (significant). Error bar represents mean ± SD. **(E)** TEM has been performed on 50nm thick sections of the high-pressure frozen COG4-mAID cells grown on sapphire discs before and after the IAA treatment (0.5 and 1 h). The scale bar is 1 µm. The red arrows indicate vesicles. **(F)** The graph represents the quantification of the total number of vesicles (50-60 nm) in the Golgi vicinity before and after IAA treatment. Two independent experiments with >10 fields were analyzed. Statistical significance was calculated by GraphPad Prism 8 using paired t-test. Here ***p ≤ 0.001 (significant). Error bar represents mean ± SD.

### Acute COG4 depletion results in rapid accumulation of COG complex-dependent (CCD) vesicles

Previous microscopy analysis of COG KO HEK293T cells revealed severe alteration of Golgi morphology, while siRNA-driven KD of COG3 and COG7 in HeLa cell also resulted in accumulation of CCD (COG Complex Dependent) vesicles (Zolov and Lupashin, 2005; Bailey Blackburn *et al*., 2016; Sumya, Pokrovskaya and Lupashin, 2021). In this study, we have employed a high-pressure freezing/freeze substitution transmission electron microscopy (TEM) approach to identify initial morphological changes in COG4-depleted RPE1 cells. TEM analysis revealed a two-fold accumulation of vesicle-like structures in a Golgi vicinity at the onset of COG4-mAID depletion (30 min after auxin addition). Most of the accumulated CCD vesicles were lacking any detectable protein coat and were situated along with the unaltered Golgi stacks. An additional two-fold increase in the number of peri-Golgi CCD vesicles was observed after one hour of IAA treatment. At this point, the Golgi stacks were moderately swollen, but the stack integrity was not severely altered (**Figure 2 E, F**). We concluded that the accumulation of CCD vesicles is the primary morphological feature of RPE1 cells acutely depleted for COG complex activity **(Figure 2E, F)**.

### Redistribution of COG sensitive Golgi v-SNAREs to CCD vesicles

COG subunits, mainly COG4, interact with the STX5-GS28-GS15-YKT6 SNARE complex to maintain intra-Golgi retrograde transport (Shestakova *et al*., 2007; Kudlyk *et al*., 2013; Laufman, Hong and Lev, 2013; Willett *et al*., 2016; Blackburn, D’Souza and Lupashin, 2019). A previous study in HeLa cells reported that COG3 KD causes the accumulation of intracellular CCD vesicles carrying Qc SNARE GS15/Bet1L and Qb SNARE GS28/GOSR1b (Zolov and Lupashin, 2005). Moreover, the GS15 and GS28 were reported as COG-sensitive GEARs-Golgi integral membrane proteins in COG mutant CHO cells (Oka *et al*., 2004). Those findings guided us to test the impact of rapid COG4 depletion on Golgi SNAREs. Golgi membranes (P30) were separated from vesicular fractions (P100) by differential centrifugation **(Figure 3A)**. Initial analysis revealed that all Golgi SNAREs were stable during the first two hours of IAA treatment **(Supplementary Figure 3A, B)** and therefore CCD vesicles were biochemically characterized after two hours of COG4-mAID depletion. As expected, WB analysis of Golgi and vesicle fractions of COG4-mAID cells revealed a significant increase in the vesicular pool of both GS15 and GS28 **(Figure 3B, C)**. Importantly, more than 70% of total cellular GS15 was relocated into CCD vesicles. At the same time, both Qa-SNARE STX5 and R-SNARE YKT6 co-fractionated with Golgi membranes upon COG4 depletion, indicating their t-SNARE role during the intra-Golgi trafficking **(Figure 3B, C)**. IF approach revealed a significant decrease in relative colocalization of GS15 and GM130 in COG depleted cells even after one hour of IAA addition **(Figure 3E, F),** while co-localization between STX5 and GM130 was not sensitive to COG4-mAID degradation **(Figure 3D, F)**. We also found that a prolonged inactivation of COG resulted in a significant decrease in the total cellular level of three Golgi SNAREs - GS15, GS28, and YKT6, indicating their dependency on the COG complex function in RPE1 cells **(Supplementary Figure 3A, 3B)**. The combined results revealed that acute COG4 depletion causes severe displacement of v-SNAREs GS15 and GS28 into relatively stable CCD vesicles.

**Figure 3.**
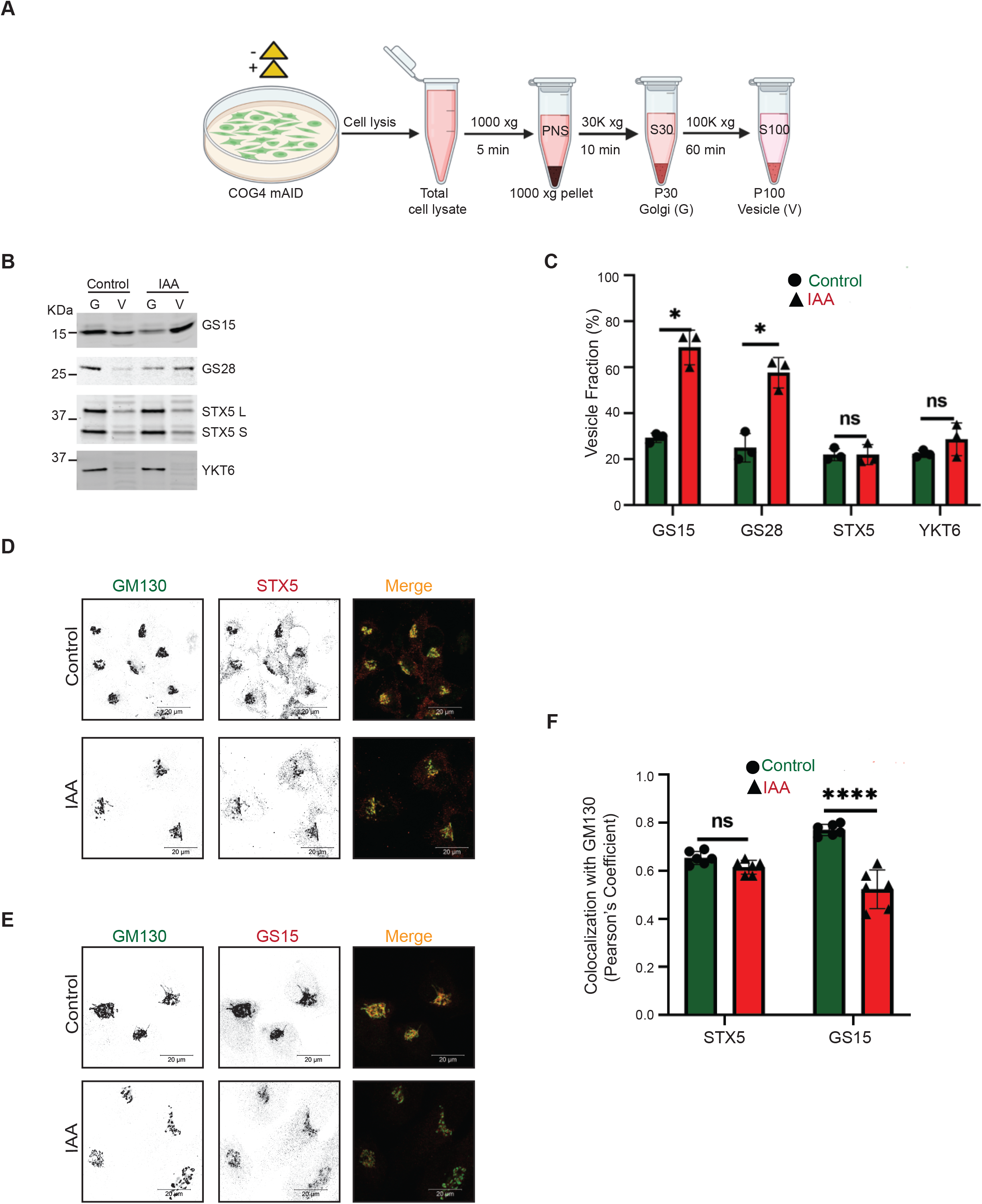
Acute COG4 depletion causes relocalization of COG-sensitive Golgi SNAREs GS15 and GS28 to the vesicular fraction. **(A)** Schematic representation of cellular fractionation experiment to prepare Golgi (G) and vesicle (V) fractions from control and IAA treated cells. **(B)** WB analysis of SNARE proteins (GS15, GS28, STX5, YKT6) in Golgi and vesicle fractions. Equal volumes of Golgi (G) and vesicle (V) membrane fractions were analyzed with antibodies as indicated. **(C)** The graph represents the quantification of vesicle fraction (%) of SNAREs in COG depleted cells compared to control. SNARE abundance in vesicles was calculated as a percentage of the fluorescent WB signal in the vesicle fraction to the combined signal in Golgi and vesicle fractions from n=3 independent experiments. Statistical significance was calculated by GraphPad Prism 8 using paired t-test. Here, p ≥ 0.05, non-significant, *p ≤ 0.01 (significant). Error bar represents mean ± SD. **(D, E)** Acute COG4 depletion does not displace the t-SNAREs (STX5) from Golgi but v-SNARE GS15 is relocalizing into vesicles. Airyscan superresolution IF analysis of untreated (control) or IAA treated COG4-mAID cells stained for **(D)** GM130 (green) and STX5 (red) and **(E)** GM130 (green) and GS15 (red). Scale bars, 20 µm. For better presentation green and red channels are shown in inverted black and white mode whereas the merged view is shown in RGB mode. **(F)** Colocalization of Golgi SNAREs with GM130 was determined by calculating Pearson’s correlation coefficient and >90 cells were analyzed. Statistical significance was calculated by GraphPad Prism 8 using paired t-test. Here, p ≥ 0.05, non-significant (ns), ****p ≤ 0.0001 (significant). Error bar represents mean ± SD.

### Differential effect of acute COG4-mAID depletion on golgins and Rab proteins

Next, we sought to dissect the effect of rapid COG4 depletion on COG interacting golgins (coiled-coil Golgi-located vesicular tethers) golgin84/GOLGA5, p115/USO1, GM130, giantin/GOLGB1 and TMF1 (Sohda *et al*., 2007, 2010; Miller *et al*., 2013; Willett *et al*., 2014; Blackburn, D’Souza and Lupashin, 2019). Previous studies reported that both golgin84 and giantin are COG sensitive “GEAR proteins”, while p115 is not sensitive to COG depletion in CHO cells (Oka *et al*., 2004). As shown above **(Figure 1C)**, GM130’s localization was not sensitive to rapid COG4-mAID depletion. Both p115 and golgin84 were also Golgi localized one hour after IAA treatment, while giantin and TMF1 were significantly mislocated from the Golgi to vesicle-like haze **(Figure 4A-D, Supplementary Figure 4A, D).** We designated golgins that remained on the Golgi upon COG depletion as target-tethers (t-tethers) and golgins that significantly relocated to CCD vesicles as vesicular tethers (v-tethers). We propose that even a partial depletion of COG4-mAID causes defect in tethering of a subset of intra-Golgi vesicles and we categorized these vesicles as “early” CCD vesicles; both giantin and TMF1 were associated with these vesicular carriers. A slightly longer (2 hours) COG4-mAID depletion resulted in significant relocalization of both golgin84 and TMF1 to a vesicular membrane fraction **(Figure 4E, F),** suggesting that golgin84 is located on the “late” CCD vesicles that are temporally distinct from the “early” giantin-containing transport intermediates. Importantly, p115, like GM130, was not sensitive to COG4 depletion, indicating that these two t-tethers operate from the Golgi side during vesicle tethering. Similarly, TGN located golgin97/GOLGA1 and golgin245/GOLGA4 were not sensitive to rapid COG depletion either (**Supplementary Figure 4B-D).**

**Figure 4.**
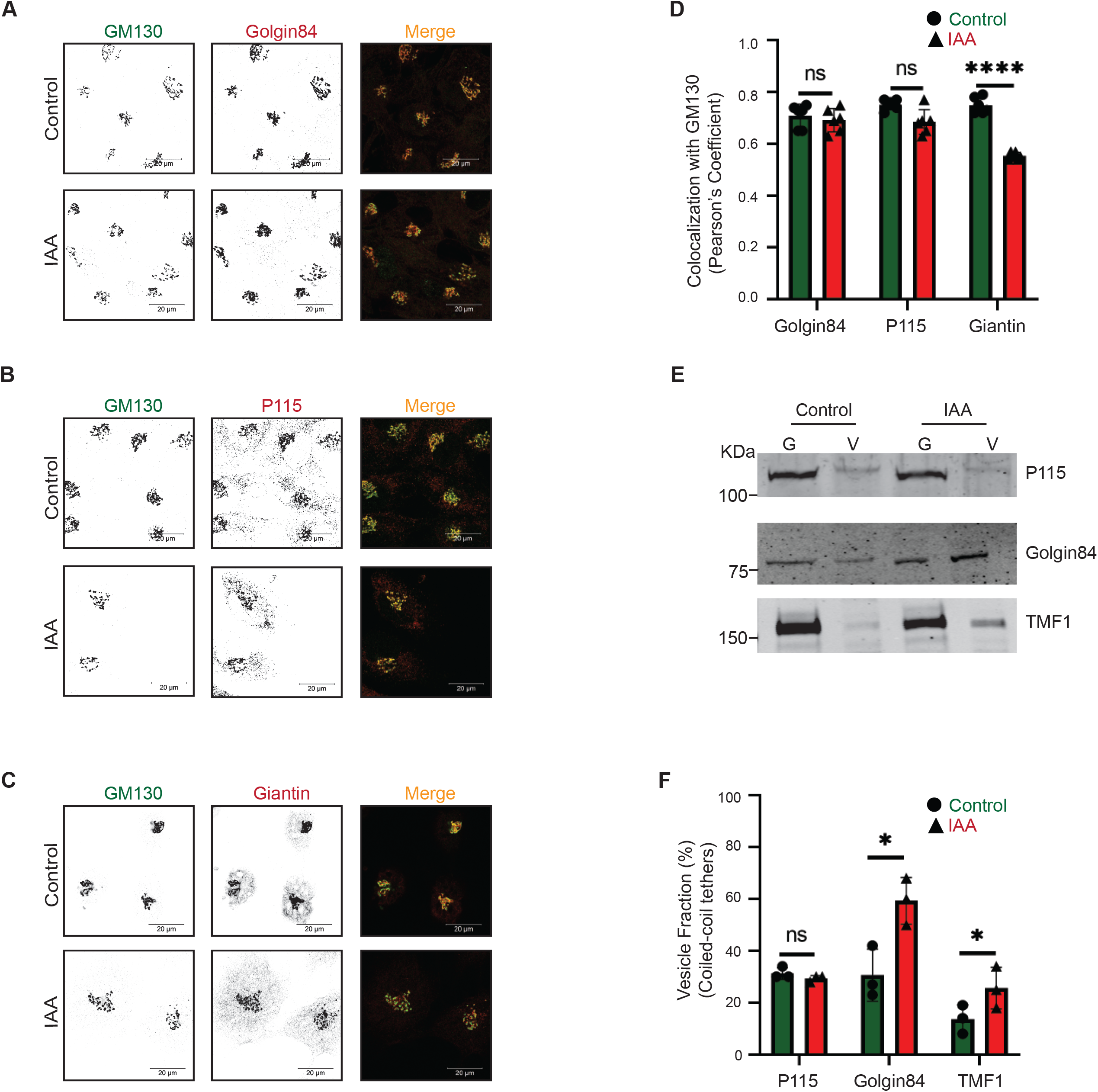
The rapid COG4 depletion has no effect on the localization of coiled-coil tether p115 but displaces giantin, golgin84, and TMF1 from Golgi. Airyscan superresolution IF analysis of untreated (control) or IAA treated COG4-mAID cells stained for **(A)** GM130 (green) and golgin84 (red), **(B)** GM130 (green) and p115 (red), and **(C)** GM130 (green) and giantin (red). Scale bars, 20 µm. For the better presentation green and red channels are shown in inverted black and white mode whereas the merged view is shown in RGB mode. **(D)** Colocalization of tested golgins with GM130 was determined by using Pearson’s correlation coefficient, >90 cells were analyzed. Statistical significance was calculated by GraphPad Prism 8 using paired t-test. Here, p ≥ 0.05, non-significant (ns), ****p ≤ 0.0001 (significant). Error bar represents mean ± SD. **(E)** WB analysis of tethers (p115, golgin84, TMF1) in Golgi and vesicle fractions. Equal volumes of Golgi (G) and vesicle (V) membrane fractions were analyzed with corresponding antibodies. **(F)** The graph represents the quantification of vesicle fraction (%) of golgins in COG-depleted cells compared to control. The abundance of p115, golgin84, and TMF1 in vesicles was calculated as a percentage of the immuno-signal in the vesicle fractions to the combined signal in Golgi and vesicle fractions from n=3 independent experiments. Statistical significance was calculated by GraphPad Prism 8 using paired t-test, *p ≤ 0.05, significant, p ≥ 0.05, non-significant (ns). Error bar represents mean ± SD.

Rabs are small GTPases involved in all steps of vesicle transport and COG interacts with several Golgi Rabs (Suvorova, Duden and Lupashin, 2002; Jaquinod *et al*., 2007; Miller *et al*., 2013). COG potentially binds to activated Rab1a, Rab1b, Rab2a, Rab4a, Rab6a, Rab10, Rab14, Rab30, Rab39, and Rab43 (Blackburn, D’Souza and Lupashin, 2019) and we have tested a representative subset of Golgi Rab proteins for their sensitivity to acute COG4-mAID depletion. Rab selection was based on the availability of commercial antibodies that can detect endogenous proteins in IF and/or WB applications. Airyscan microscopy revealed that Rab1B and Rab6A did not change their localization from the Golgi into CCD vesicles **(Figure 5A-C)** after one hour of COG depletion. Similar results were obtained with cells transiently transfected with GFP-Rab30a and GFP-Rab43a **(Supplementary Figure 5A-C)**. Two hours of COG4-mAID depletion did not shift Rab2A, Rab6A, and Rab30 into CCD vesicles **(Figure 5D, E)**. By contrast, Rab1B vesicular fraction was increased significantly at this time point, indicating that Rab1B could be incorporated into “late” CCD vesicles **(Figure 5D, E)**. In summary, COG depletion did not shift the majority of COG-interacting Rabs to the vesicular fraction, strongly suggesting that these Rabs, like t-tethers p115 and GM130, primarily operate from the Golgi side during the vesicle tethering process.

**Figure 5.**
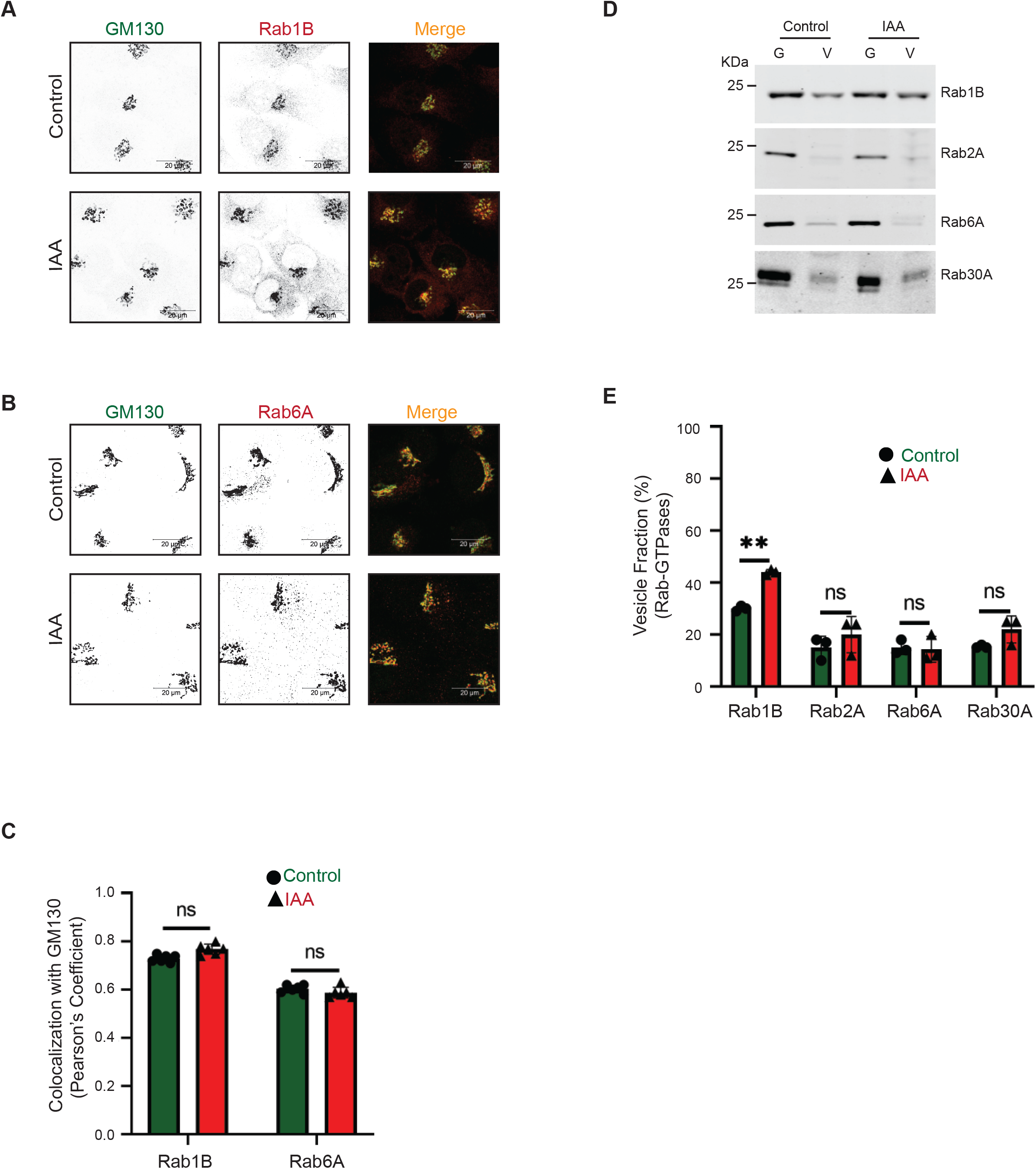
The majority of COG-interacting Golgi Rabs are not changing their localization upon COG4 depletion. Airyscan superresolution IF analysis of untreated (control) or IAA treated COG4-mAID cells stained for **(A)** GM130 (green) and Rab1B (red), and **(B)** GM130 (green) and Rab6 (red). Scale bars, 20 µm. For better presentation green and red channels are shown in inverted black and white mode whereas the merged view is shown in RGB mode. **(C)** Colocalization of Rab-GTPases with GM130 was performed by calculating Pearson’s correlation coefficient, >90 cells were analyzed. Statistical significance was calculated by GraphPad Prism 8 using paired t-test. Here, p ≥ 0.05, non-significant (ns). Error bar represents mean ± SD. **(D)** WB analysis of Rab1B, Rab2A, Rab6A, and Rab30A in Golgi and vesicle fractions. Equal volumes of Golgi (G) and vesicle (V) membrane fractions were analyzed with antibodies as indicated. **(E)** The graph represents the quantification of vesicle fraction (%) of Rab-GTPases in COG depleted cells compared to control. Rab-GTPase abundance in vesicles was calculated as a percentage of the immuno signal in the vesicle fraction to the combined signal in Golgi and vesicle fractions from n=3 independent experiments. Statistical significance was calculated by GraphPad Prism 8 using paired t-test. Here, p ≥ 0.05, non-significant and **p ≤ 0.01. Error bar represents mean ± SD.

### Redistribution of Golgi resident proteins into CCD vesicles

One of the most common defects associated with permanent COG dysfunction is the defect in the stability of intracellular recycling glycoproteins and Golgi enzymes (Oka *et al*., 2004; I. Pokrovskaya *et al*., 2011; Blackburn *et al*., 2018; Blackburn, D’Souza and Lupashin, 2019; Khakurel *et al*., 2021; Sumya, Pokrovskaya and Lupashin, 2021). COG-related depletion of Golgi proteins could be caused either by direct re-routing of recycling protein to degradative compartments like lysosomes, or their instability could be due to deficient glycosylation, or their relocalization into recycling CCD vesicles that are unable to dock and fuse to a proper compartment and therefore got degraded by yet unknown mechanism (Oka *et al*., 2004; Zolov and Lupashin, 2005; Shestakova, Zolov and Lupashin, 2006; I. Pokrovskaya *et al*., 2011; Bailey Blackburn *et al*., 2016; Blackburn *et al*., 2018; Blackburn, D’Souza and Lupashin, 2019; Khakurel *et al*., 2021; Sumya, Pokrovskaya and Lupashin, 2021). To identify the primary defect associated with COG dysfunction, we first determined changes in the total cellular level of Golgi recycling and resident proteins upon both acute and prolonged COG4-mAID depletion **(Figure 6A, B, 7A, B)**. Results revealed that cis-Golgi GPP130/GOLIM4, medial-Golgi TMEM165, trans-Golgi TGN46/TGOLN2, and SDF4/Cab45 were stable during the first two hours of IAA treatment and then degraded during a prolonged COG depletion. In the case of TGN46 and SDF4, protein depletion coincided with a change in protein electrophoretic mobility, indicating defects in secondary protein modifications **(Figure 6A, B)**. A similar degradative pattern was observed for Golgi enzymes B4GalT1 (beta-1, 4-galactosyltransferase 1), GalNT2 (N-acetylgalactosaminyltransferase 2), and MGAT1 (alpha-1,3-mannosyl-glycoprotein 2-beta-N-acetylglucosaminyltransferase 1) (**Figure 7A, B)**, indicating that only prolonged, but not acute depletion of COG4-mAID causes degradation of Golgi resident proteins. At the same time, Airyscan microscopy analysis revealed a significant fraction of all tested Golgi resident proteins, with the notable exception of GalNT2, displaced from the Golgi into a vesicle-like dot pattern (**Figure 6C-G, 7C, D, F)** within one hour of COG4 depletion.

**Figure 6.**
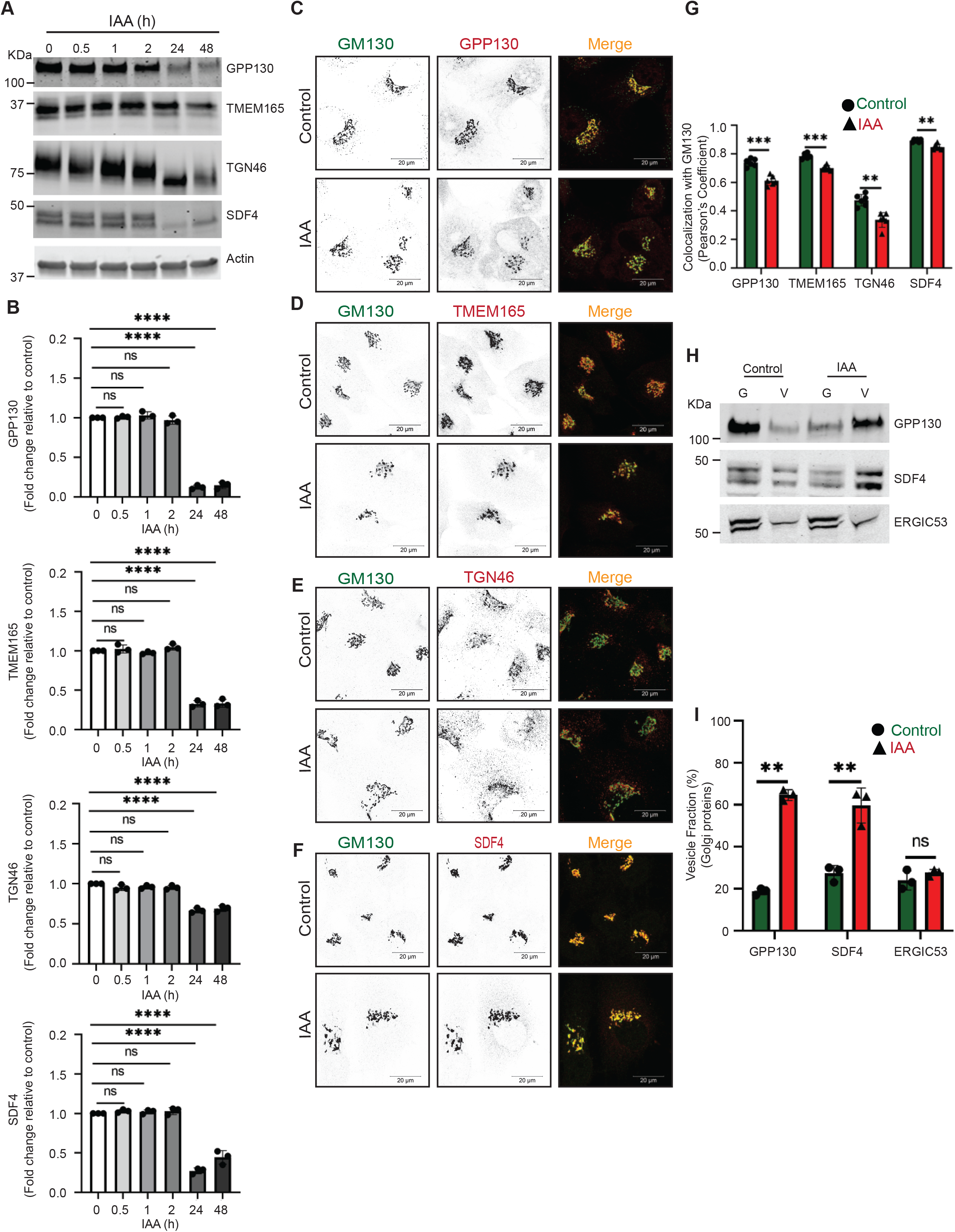
Acute COG4 depletion causes relocalization of Golgi resident proteins into CCD vesicles. **(A)** WB of time-dependent depletion of COG4-mAID shows the expression of Golgi resident proteins (GPP130, TMEM165, TGN46, and SDF4). 10 µg of total cell lysates were loaded and probed with indicated antibodies. β actin was used as a loading control. **(B)** The graph represents the quantification of **A**. In the bar graph, values represent the mean ± SD from three independent experiments. Statistical significance was calculated using one-way ANOVA. P≥0.05, non-significant (ns), ****p ≤ 0.0001, significant. **(C, D, E, F)** Airyscan superresolution IF analysis of untreated (control) or auxin treated (IAA) COG4-mAID cells stained for **(C)** GM130 (green) and GPP130 (red), **(D)** GM130 (green) and TMEM165 (red), **(E)** GM130 (green) and TGN46 (red), **(F)** GM130 (green) and SDF4 (red) respectively. Scale bars, 20 µm. For better presentation green and red channels are shown in inverted black and white mode whereas the merged view is shown in RGB mode. **(G)** Colocalization of Golgi resident proteins with GM130 was determined by calculating Pearson’s correlation coefficient and >90 cells were analyzed. Statistical significance was calculated by GraphPad Prism 8 using paired t-test. Here, ****p ≤ 0.0001, ***p ≤ 0.001, *p ≤ 0.05, significant and p ≥ 0.05 non-significant (ns). Error bar represents mean ± SD. **(H)** WB analysis of GPP130, SDF4, and ERGIC53 in Golgi and vesicle fractions. Equal volumes of Golgi (G) and vesicle (V) membrane fractions were analyzed with antibodies as indicated. **(I)** The graph represents the quantification of vesicle fraction (%) of Golgi resident proteins in COG depleted cells compared to control. The Golgi protein abundance in vesicles was calculated as a percentage of the immuno signal in the vesicle fraction to the combined signal in Golgi and vesicle fractions from n=3 independent experiments. Statistical significance was calculated by GraphPad Prism 8 using paired t-test, ****p ≤ 0.0001, ***p ≤ 0.001, *p ≤ 0.05, significant and p ≥ 0.05 non-significant (ns). Error bar represents mean ± SD.

**Figure 7.**
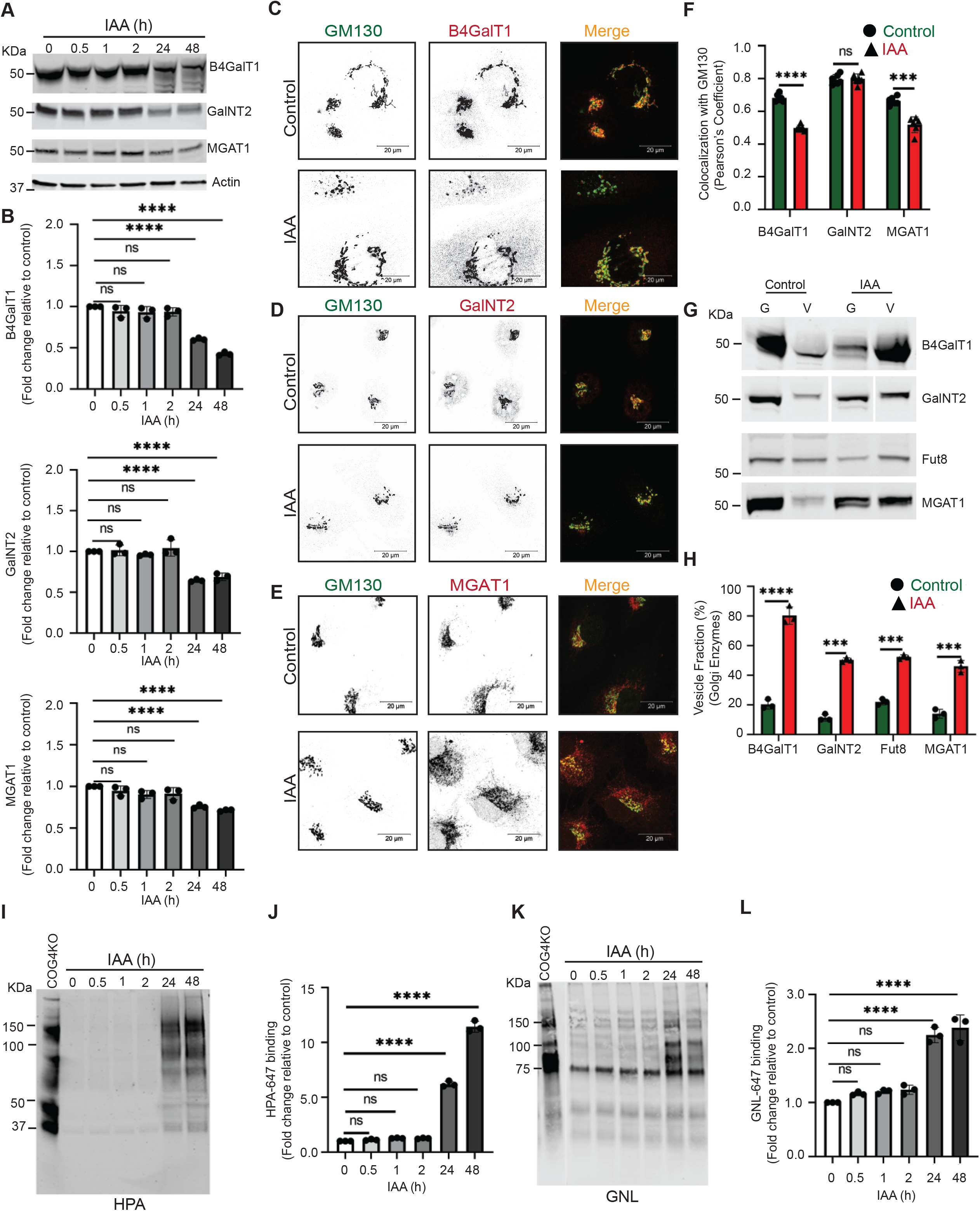
Acute depletion of COG4 displaces a significant fraction of the Golgi enzymes into CCD vesicles causing their subsequent degradation and glycosylation defects. **(A)** WB of total cell lysates during time-dependent depletion of COG4-mAID shows the expression of Golgi B4GalT1, GalNT2, and MGAT1 enzymes. 10 µg of total cell lysates were loaded and probed with indicated antibodies. β actin was used as a loading control. **(B)** The graph represents the quantification of **A**. In the bar graph, values represent the mean ± SD from three independent experiments. Statistical significance was calculated using one-way ANOVA. ****p ≤ 0.0001, significant. **(C, D, E)** Airyscan superresolution IF analysis of untreated (control) or IAA treated COG4-mAID cells stained for **(C)** GM130 (green) and B4GalT1 (red), **(D)** GM130 (green) and GalNT2 (red), **(E)** GM130 (green) and MGAT1 (red) respectively. Scale bars, 20 µm. For better presentation green and red channels are shown in inverted black and white mode whereas the merged view is shown in RGB mode. **(F)** Colocalization of Golgi enzymes and GM130 was determined by calculating Pearson’s correlation coefficient, >90 cells were analyzed. Statistical significance was calculated by GraphPad Prism 8 using paired t-test. Here, ***p ≤ 0.001, ****p ≤ 0.0001 (significant) and p ≥ 0.05 non-significant (ns). Error bar represents mean ± SD. **(G)** WB analysis of glycosylation enzymes in Golgi and vesicle fractions. Equal volumes of Golgi (G) and vesicle (V) membrane fractions were analyzed with corresponding antibodies. **(H)** The graph represents the quantification of vesicle fraction (%) of Golgi enzymes in COG depleted cells compared to control. The Golgi enzyme abundance in vesicles was calculated as a percentage of the immuno signal in the vesicle fractions to the combined signal in Golgi and vesicle fractions from n=3 independent experiments. Statistical significance was calculated by GraphPad Prism 8 using paired t-test, ****p ≤ 0.0001, ***p ≤ 0.001, significant. Error bar represents mean ± SD. **(I)** HPA-647 lectin blot of total cell lysates obtained during time-dependent depletion of COG4-mAID. COG4 KO cells were used as a control. **(J)** The graph represents the quantification of **H. (K)** GNL-647 lectin blot of total cell lysates during time-dependent depletion of COG4-mAID. COG4 KO cells were used as a control. **(L)** The graph represents the quantification of **K.** Values in bar graphs represent the mean ± SD from three independent experiments. Statistical significance was calculated in GraphPad Prism 8 using one-way ANOVA. ***p ≤ 0.001, **p ≤ 0.01, *p ≤ 0.05.

To further investigate the COG-dependent behavior of Golgi resident proteins, we analyzed Golgi and vesicle fractions as described above **(Figure 3A)**. WB analysis revealed that the vesicular pool of all tested Golgi recycling proteins and enzymes including trans-Golgi enzyme FUT8 (Alpha-(1,6)-fucosyltransferase 8) increased significantly after two hours of COG4 depletion **(Figure 6H-I, 7G-H)**. COG depletion resulted in the relocalization of ∼ 50% of all analyzed Golgi resident proteins into CCD vesicles. Most dramatic relocalization (∼80%) to the vesicular fraction was observed for B4GalT1, indicating that this trans-Golgi enzyme is constantly recycling in CCD vesicles. Interestingly, GalNT2 partially relocalized to a vesicular fraction as well, suggesting that this enzyme is mostly recycled by the “late” CCD vesicles. The relocation of Golgi resident proteins to vesicular fraction was specific since the localization of another recycling protein ERGIC53/LMAN1 did not change its distribution between Golgi and vesicle fractions upon COG4-mAID depletion **(Figure 6H-I)**.

Fluorescently-labeled lectins are a useful tool to assess the functionality of glycosylation machinery (Dan, Liu and Ng, 2016). Others and we reported altered binding of several lectins to both total and surface-exposed glycoprotein in COG mutants due to impaired Golgi glycosylation (Shestakova, Zolov and Lupashin, 2006; Richardson *et al*., 2009; I. Pokrovskaya *et al*., 2011; Peanne *et al*., 2011; Reynders *et al*., 2011; Bailey Blackburn *et al*., 2016; Khakurel *et al*., 2021; Sumya, Pokrovskaya and Lupashin, 2021). Galanthus nivalus lectin (GNL) binds to terminal 1,3- and 1,6-linked mannose residues on N-linked glycans (Kaku and Goldstein, 1992; Bailey Blackburn *et al*., 2016; Khakurel *et al*., 2021; Sumya, Pokrovskaya and Lupashin, 2021), while Helix Pomatia Agglutinin (HPA) binds to terminal N-acetylgalactosaminyl residues in O-glycans (Brooks, 2000; Khakurel *et al*., 2021; Sumya, Pokrovskaya and Lupashin, 2021) respectively. Therefore, an increase in binding of GNL indicates MGAT1 deficiency and accumulation of underglycosylated N-linked glycoconjugates, while an increased binding of HPA indicates a deficiency in GalNT enzymes and accumulation of underglycosylated O-linked glycoconjugates respectively (Brooks, 2000; Irina Pokrovskaya *et al*., 2011; Khakurel *et al*., 2021; Sumya, Pokrovskaya and Lupashin, 2021). Both MGAT1 and GalNT2 are severely mislocated from the Golgi after the acute COG depletion. To check the ‘N’ and ‘O’-glycosylation fidelity during time-dependent COG4 depletion, GNL and HPA conjugated with Alexa-647 (GNL-647 and HPA647) have been utilized for WB analysis of COG4-mAID cells **(Figure 7I-L)**. WB lectin analysis revealed that GNL-647 and HPA-647 binding was not increased during the first two hours of COG depletion, indicating a lack of detectable glycosylation defects. This data indicated that previously observed changes in glycosylation in COG mutant cells are a secondary manifestation of COG complex deficiency. In agreement with this hypothesis, GNL and HPA binding were significantly increased after prolonged COG4 depletion **(Figure 7I-L)**. In combination, our data revealed that the acute COG4 depletion causes the redistribution of the Golgi enzymes and a subset of other resident proteins into CCD vesicles whereas a prolonged COG depletion causes significant degradation of mislocated enzymes causing glycosylation defects.

### COG sensitive Golgi proteins are recycled in distinct CCD vesicles

Previously, we showed that COG3 KD in HeLa cells resulted in relatively slow (48-72 hours) accumulation of several Golgi proteins in CCD vesicles (Zolov and Lupashin, 2005). The acute COG4-mAID depletion approach resulted in a much faster (1-2 hours) accumulation of CCD vesicles containing a subset of COG interacting partners, Golgi enzymes as well as Golgi resident proteins. IF and WB data indicate that several proteins (like GS15, MGAT1, and B4GalT1) dislocate from the Golgi at the onset of COG4-mAID depletion in the “early” CCD vesicles, while others (like Rab1B and GalNT2) require a longer depletion of COG for their relocalization into “late” CCDs. Temporal differences in the displacement of different Golgi proteins into vesicular fractions indicate that different resident proteins are using different carriers for recycling between Golgi subcompartments. To test this hypothesis, we first analyzed CCD vesicles by Airyscan microscopy. In control cells, MGAT1 and B4GalT1 reside in adjoined Golgi cisternae, showing partial colocalization by IF **(Figure 8A)**. If these two proteins use the same CCD vesicles for their COG-dependent recycling, we expect an increase in their colocalization, since vesicle size (∼60 nm) is significantly below the resolution limits of Airyscan microscopy (170 nm). Therefore, if two different proteins reside in the same vesicle, they would show strong colocalization. Indeed, if two different secondary antibodies labeled with Alexa488 and Alexa647 were used to analyze B4GalT1 localization, significant colocalization of fluorescent signals was observed (F.T.S. unpublished data). In contrast, Airyscan analysis revealed a significant decrease in colocalization of MGAT1 and B4GalT1 upon acute COG4-mAID depletion, indicating that these two proteins recycle in different sets of CCD vesicles **(Figure 8B)**. Similar results have been obtained for B4GalT1 and cis-Golgi protein GPP130 indicating that these proteins also travel in separate vesicles **(Supplementary Figure 6A, B)**. To complement IF studies, the analysis of CCD vesicles separated via sucrose velocity sedimentation (Love *et al*., 1998) was performed **(Figure 8C)**. The WB analysis of individual sucrose gradient fractions revealed vesicular fractions enriched for B4GalT1, MGAT1, and GalNT2 proteins **(Figure 8D)**. The maximum signal for B4GalT1 was detected in fraction 5 while MGAT1 and GalNT2 are mostly enriched in fraction 6 **(Figure 8E)** indicating that medial and trans-Golgi enzymes are recycling in separate CCD vesicle populations that slightly differ from each other by size and/or density. WB analysis of gradient fractions for B4GalT1 and GPP130 **(Supplementary Figure 6C)** also revealed two distinct vesicular populations **(Supplementary Figure 6D)**. In summary, the IF and WB analysis of vesicles accumulated in acute COG depleted cells indicate that cis, medial and trans-Golgi residents recycle in different populations of CCD vesicles.

**Figure 8:**
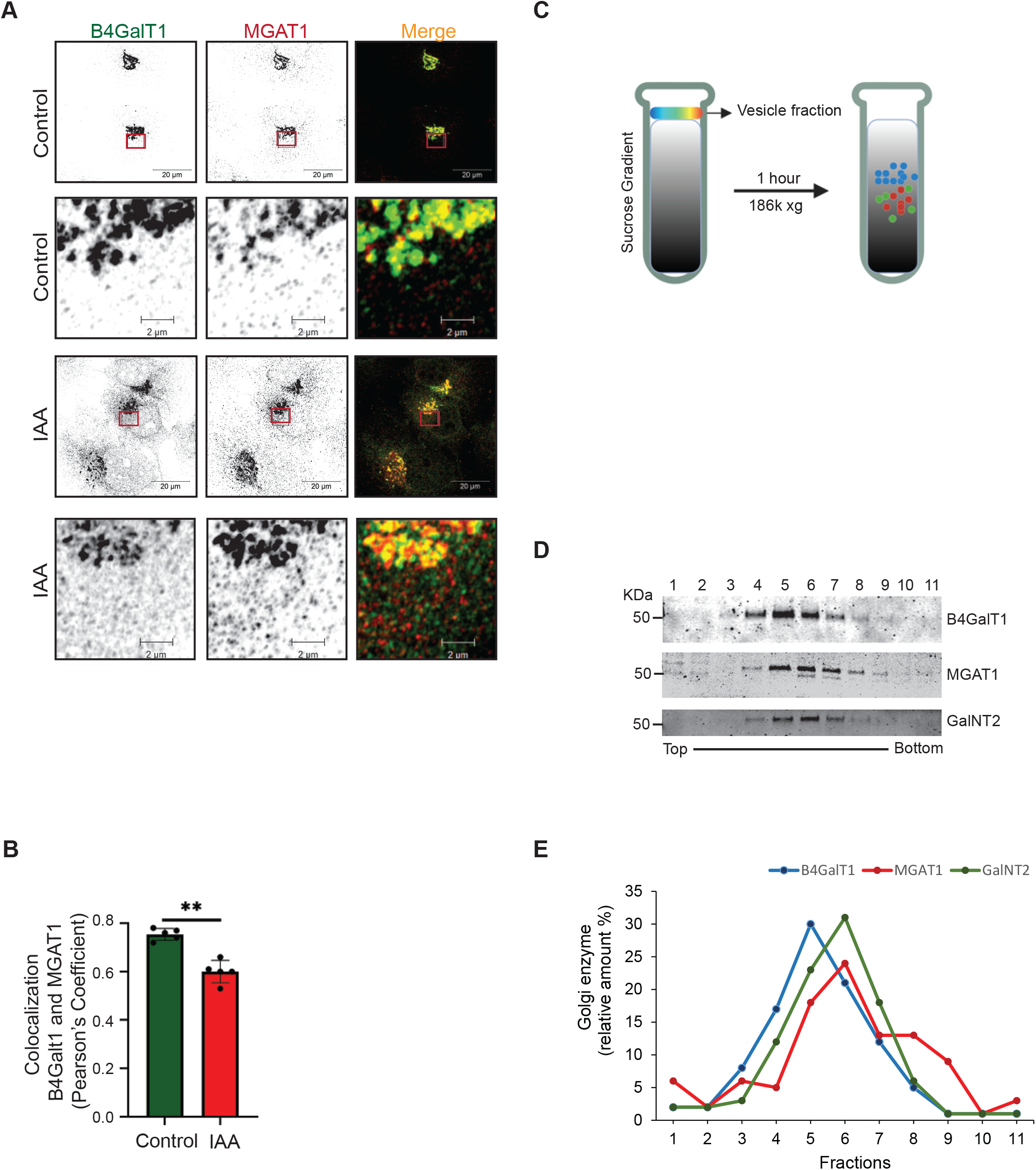
B4GalT1 and MGAT1 recycle in distinct CCD vesicles. **(A)** Airyscan superresolution IF analysis of untreated (control) or IAA treated COG4-mAID cells stained for B4GalT1 (green) and MGAT1 (red). Scale bars, 20 µm. The enlarged (5x) view of the framed area of the upper panel is shown in the bottom panel. For better presentation green and red channels are shown in inverted black and white mode whereas the merged view is shown in RGB mode. **(B)** Colocalization of B4GalT1 with MGAT1 was determined by calculating Pearson’s correlation coefficient, >90 cells were analyzed. Statistical significance was calculated by GraphPad Prism 8 using paired t-test. Here, **p ≤ 0.001, (significant). Error bar represents mean ± SD. **(C)** Schematic representation of CCD vesicle fractionation by sucrose velocity gradient centrifugation. **(D)** WB of vesicle fractions separated by sucrose gradient fractions tested for B4GalT1, MGAT1, and GalNT2. **(E)** The line graph represents the quantification of **D.**

### Displacement of multiple vesicular coats from Golgi upon acute COG4 depletion

It has been postulated that intra-Golgi retrograde trafficking is primarily accomplished by selective incorporation of recycling proteins into vesicles formed by COPI protein coat (Béthune *et al*., 2006; Popoff *et al*., 2011; Luo and Boyce, 2019). COG-COPI genetic and physical interaction was reported by others and us (Ram, Li and Kaiser, 2002; Suvorova, Duden and Lupashin, 2002; Oka *et al*., 2004; Zolov and Lupashin, 2005; Willett, Ungar and Lupashin, 2013), suggesting that COG is tethering COPI-formed intra-Golgi trafficking intermediates. At the same time, recent data from the yeast system demonstrated the role of the AP1 vesicle coat complex in recycling trans-Golgi enzymes (Gu, Crump and Thomas, 2001; Casler *et al*., 2019; Duncan, 2022). Also, GGA vesicular coat was implicated in the localization of a subset of Golgi proteins (Costaguta *et al*., 2001; Popoff *et al*., 2011).

Since our analysis revealed that different resident Golgi proteins are recycling in different CCD vesicles we have investigated the changes in the localization of Golgi located vesicular coat machineries in cells acutely depleted for the COG complex. Firstly, the localization of β and γ subunits of COPI vesicular coat complex was determined by Airyscan IF approach. Microscopy analysis of COG4-mAID cells revealed that both β COP/COPB2 and γCOP/COPG1 were Golgi located in control cells and become severely displaced from Golgi perinuclear region as early as the one hour of COG4 depletion **(Figure 9A, 9B)**. COPI selects recycling proteins into vesicles either directly (Liu, Doray and Kornfeld, 2018), or indirectly, using additional adaptors like GOLPH3 and GOLPH3L (Schmitz *et al*., 2008; Tu *et al*., 2008; Welch *et al*., 2021). In agreement with this model, GOLPH3 was rapidly displaced from the Golgi region at the early onset of COG4-mAID depletion (**Supplementary Figure 7A, B).** Biochemical fractionation of COG depleted cells did not reveal COPI associated with CCD vesicle fraction, indicating that upon COG4-mAID depletion COPI is dissociated from non-tethered vesicles and accumulated in cytosol (F.T.S. unpublished data). Intriguingly, two other Golgi-based vesicle adaptor complexes, GGA and AP1 reacted to COG acute repletion in a manner similar to COPI. The GGAs and the AP1 are mainly localized in the trans-Golgi network and are likely to select cargo proteins into distinct types of vesicles (Robinson, 2004). IF analysis showed that both GGA2 and AP1β/AP1B1 adaptor proteins were displaced from Golgi upon acute COG depletion **(Figure 9C, 9D)**. A colocalization analysis confirmed significant decrease in colocalization of COPI subunits (β□, γ) as well as adaptor proteins (GGA2, AP1β) with the Golgi marker GM130 **(Figure 9E)**. In summary, we have uncovered that COG complex acute depletion caused a buildup of multiple types of non-tethered recycling vesicles that likely to be formed from different Golgi cisternae with the help of COPI, AP1, and GGA vesicle budding/cargo sorting machineries.

**Figure 9.**
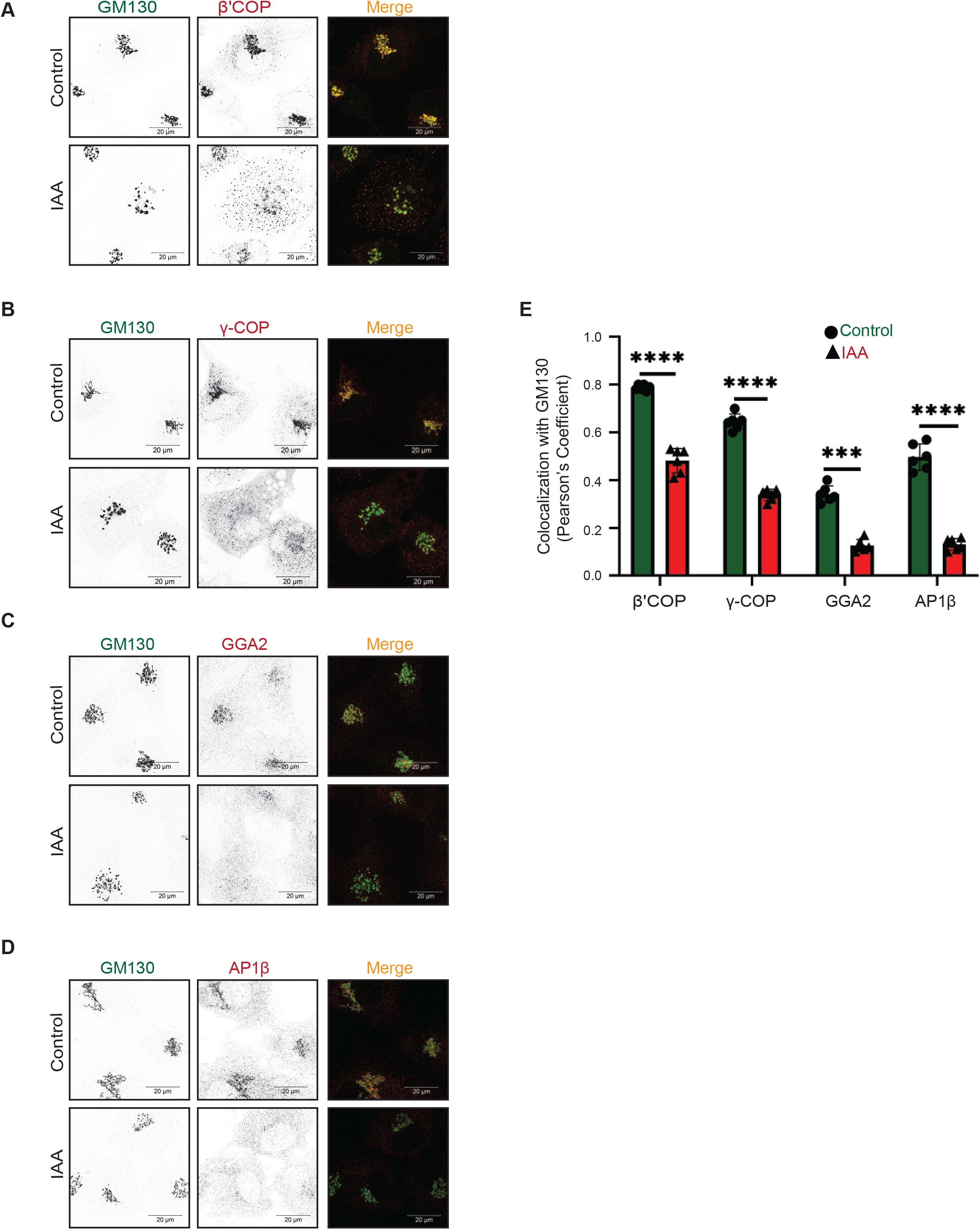
Acute COG4 depletion displaces multiple vesicular coats from Golgi. **(A)** Airyscan superresolution IF analysis of untreated (control) or IAA treated) COG4-mAID cells stained for GM130 (green) and β COP (red), **(B)** GM130 (green) and γCOP (red), **(C)** GM130 (green) and GGA2 (red), **(D)** GM130 (green) vs AP1-β (red) respectively. Scale bars, 20 µm. For the better presentation green and red channels are shown in inverted black and white mode whereas the merged view is shown in RGB mode. **(E)** Colocalization of coat proteins with GM130 was determined by calculating Pearson’s correlation coefficient, >90 cells were analyzed. Statistical significance was calculated by GraphPad Prism 8 using paired t-test. Here, ***p ≤ 0.001 (significant) and ****p ≤ 0.0001 (significant). Error bar represents mean ± SD.

## Discussion

Intra-Golgi trafficking and localization of Golgi resident proteins are intensely studied for more than 50 years, but the exact rules for localization of Golgi glycosylation machinery, the repertoire of membrane carriers that move retrograde and anterograde cargo, and the mechanisms for vesicle tethering and docking in the Golgi are still an enigma. To advance our knowledge of Golgi physiology, we investigated the effect of the acute depletion of the COG complex, the major multisubunit Golgi vesicle tethering factor, on the dynamics of Golgi resident proteins. We found that rapid COG4 depletion was sufficient to induce COG complex dysfunction causing a dramatic accumulation of CCD vesicles that carry a specific set of SNAREs, golgins, and Golgi resident proteins. In agreement with previous EM data (Orci *et al*., 1997), accumulated intra-Golgi vesicles were mostly uncoated and initially located in close proximity to the unaltered Golgi stack, indicating that the coat falls off promptly after vesicle budding to allow golgin-assisted long-range tethering to occur. A large group of Golgi resident proteins promptly relocated into CCD vesicles 30-60 min after the initiation of COG4 depletion, therefore, we termed these carriers the “early” CCD vesicles. Other Golgi residents required a more substantial two-hour-long COG depletion for their incorporation into vesicular carriers suggesting that these molecules travel in the “late” CCD vesicles.

In agreement with previously published data, we found that GS15 and GS28 are actively incorporated into CCD vesicles to operate as v-SNARE proteins, while STX5 and Ykt6 remain on the Golgi and work as t-SNAREs (Zhang and Hong, 2001). STX5 is a transmembrane protein, which is shown to cycle via ER (Rowe *et al*., 1998; Bentley *et al*., 2006; Cottam and Ungar, 2012; Linders *et al*., 2020); it will be important to investigate which membrane carriers are used to recycle STX5 during Golgi biogenesis (Rowe *et al*., 1998). Curiously, previous proteomic studies identified STX5 as a component of *in vitro* formed COPI Golgi vesicles (Cottam and Ungar, 2012; Adolf *et al*., 2019; Linders *et al*., 2019) this result is likely to indicate the principal difference between *in vivo* accumulated and *in vitro* formed Golgi-derived vesicles. YKT6 is anchored to the membrane by prenylation, which is essential for Golgi integrity and cell viability (PMID: 33035318, PMID: 32128853), further supporting Ykt6 role as a t-SNARE at the Golgi.

Rab GTPases are traditionally described as a vesicular component of trafficking machinery (Li and Marlin, 2015; Jin *et al*., 2021). Rab1, Rab2, Rab6, and Rab30 proteins are incorporated in the *in vitro-*produced Golgi-derived COPI vesicles (Gilchrist *et al*., 2006; Bhuin and Roy, 2014; Adolf *et al*., 2019). Surprisingly, our data revealed that many tested Golgi Rab proteins (Rab2a, Rab6, Rab30, Rab43) did not change their intracellular localization upon the acute COG depletion and remained on the Golgi membranes. This data again indicated that the *in vivo* formed and physiologically relevant CCD vesicles are significantly distinct from the *in vitro*-produced COPI vesicular structures. One possibility is that the *in vitro* formed COPI vesicles are lacking regulatory machinery that limits the incorporation of STX5 and Rabs and favors the concentration of Golgi resident proteins into vesicular carriers. As Rab2, Rab6, Rab30, and Rab43 interact with the COG complex (Willett *et al*., 2014; Blackburn, D’Souza and Lupashin, 2019) we propose that they preferentially work from the Golgi membrane to tether incoming CCD vesicles. In agreement with this proposal, yeast Golgi Rab Ypt1, which is highly related to Rab2, Rab30, and Rab43 (Liu and Storrie, 2012) is shown to work from the Golgi membrane during tethering of COPII vesicles (Cao and Barlowe, 2000). Unlike other tested Rabs, Rab1B was partially relocated to the “late” CCD vesicles, and therefore could work from the vesicle side during the tethering/fusion process.

Coiled-coil Golgi-located vesicular tethers, golgins, like SNAREs, demonstrated differential response to the acute COG depletion. Two COG-interacting golgins, p115 and GM130 remained on the Golgi and are likely to operate in vesicle tethering from the Golgi side as t-tethers. In contrast, giantin, golgin84, and TMF1 were actively relocated to CCD vesicles, indicating that these golgins may operate from the vesicle side as v-tethers. Both giantin and golgin84 are transmembrane proteins, therefore, their segregation into CCD vesicles could not only indicate their function as v-tethers but also their behavior as a recycling Golgi cargo. On the other hand, TMF1 protein does not have a transmembrane domain and must be actively segregated to CCD vesicles. Previous *in vitro* studies indicated that during tethering of intra-Golgi vesicles golgin84 interacts with the CASP, while giantin interacts with p115 (Malsam *et al*., 2005). Our results showed that one component (golgin84 or giantin) of this tethering reaction is vesicle localized, while the other component (p115 and possibly CASP) is associated with the Golgi stack. It is also interesting to note that previous data implicated Rab1, Rab2a, and Rab6 as TMF1 binding partners (Fridmann-Sirkis, Siniossoglou and Pelham, 2004; Miller *et al*., 2013). In COG-depleted cells, Rab2a and Rab6 remained associated with the Golgi, while TMF1 and Rab1b are associated with CCD vesicles, suggesting that TMF1 initiates vesicle tethering by binding to activated Rabs on both vesicle and Golgi membranes.

The most intriguing aspect of our work is finding that upon the acute COG depletion the majority of Golgi resident proteins are significantly relocated into CCD vesicles and that these vesicles are not uniform in their content and physical properties. Massive accumulation of Golgi resident proteins in Golgi-derived vesicles could be viewed as a surprising result, considering that previous immuno-EM analysis failed to localize Golgi enzymes in budding profiles and peri-Golgi vesicles (Orci, Amherdt, *et al*., 2000; Kweon *et al*., 2004). This may be due to the very short half-life of intra-Golgi vesicles in cells with functional COG complex and with a small number of recycling enzymes carried by one transport vesicle. It is likely that CCD vesicles originated from different Golgi subcompartments and that different vesicle budding machinery are involved in their formation. Some Golgi residents, like medial-Golgi MGAT1 and trans-Golgi B4GalT1, are very sensitive to COG depletion, actively relocating to the “early” CCDs, indicating that both enzymes are frequently incorporated into transport membrane intermediates that require COG to tether and fuse with the acceptor membrane. Importantly, we were unable to colocalize MGAT1 and B4GalT1 in the same peri-Golgi vesicles, indicating that these proteins recycle in different populations of “early” CCD vesicles. The difference in rates at which different Golgi residents are incorporated into CCD vesicles could be related to the precision of their localization in specific Golgi cisternae. MGAT1 and B4GalT1 operate in partially overlapping medium and trans-Golgi cisternae (Nilsson *et al*., 1993; Rabouille *et al*., 1995). To maintain this specific localization in a constantly maturing Golgi, these enzymes should be incorporated into the recycling CCD vesicles at a higher rate. On the other hand, GalNT2 is found throughout the Golgi stack (Röttger *et al*., 1998). This type of localization requires less precision and therefore GalNT2 could be recycled from multiple Golgi compartments and incorporated into CCD vesicles at a lesser rate. Indeed, we found that GalNT2 is less sensitive to COG depletion, relocating to “late” COG vesicles. V-tether golgin84 is also reacted “late” to COG depletion, suggesting that this protein was using GalNT2-filled carriers. However, the mitochondria relocalization studies demonstrated that ectopically expressed golgin84 tethers membranes containing GalNT2 (Wong and Munro, 2014), indicating that golgin84 and GalNT2 used different CCD vesicles for their recycling of needs. In agreement with IF data, vesicles carrying medial and trans-Golgi enzymes were separated on a velocity gradient. Similar separation of vesicles carrying MGAT1 and B4GalT1 was observed previously by the Ostermann laboratory while studying the *vitro* formed COPI Golgi vesicles (Love *et al*., 1998), suggesting that the “early” CCD vesicles accumulated in COG deficient cells are formed by COPI machinery. Indeed, several COPI subunits and COPI-interacting adaptor GOLPH3 were very sensitive to COG depletion. Surprisingly, two other Golgi-located vesicular coats AP1 and GGA, were also sensitive to acute COG4 depletion, indicating that CCD vesicles could be formed by at least three different vesicular coats. The involvement of the AP1 complex in the intra-Golgi trafficking was recently demonstrated in yeast cells by the Glick laboratory (Casler *et al*., 2019) and our findings support the notion that AP1 is playing an active role in intra-Golgi trafficking in mammalian cells as well. The role of the GGA coat in the localization of Golgi enzymes has been documented previously (Daboussi *et al*., 2017).

In a summary model **(Figure 10)** we postulate that during the Golgi maturation process all Golgi resident proteins are continuously recycling from the later cisternae to the early ones in CCD vesicles. The notable exception from this rule is STX5, which is not incorporated into CCD carriers and is likely to be recycled by tubular connections or other COG-independent mechanisms. The likelihood of protein incorporation into CCD vesicle and the rate of vesicular recycling is different for different Golgi residents and it may define their intra-Golgi localization pattern – residents with tight localization (like B4GalT1) would cycle more frequently compared to more dispersed residents like GalNT2. CCD carriers are formed from different Golgi cisternae by specific coat machinery, such as COPI is forming vesicles from cis, medial, and trans compartments, while AP1 is selecting resident proteins in trans-Golgi and TGN, and GGA is forming retrograde vesicles from TGN and the endosomal compartments. Different CCD vesicles carry distinct cargo and a specific set of v-tethers. It is currently unknown how many molecules of Golgi residents are packaged in a single 60 nm vesicle carrier, but our inability to colocalize Golgi enzymes in the same CCD vesicle suggests a relatively low number of cargo molecules per vesicle. GS28 and GS15 are likely to be the major v-SNAREs for all classes of CCD vesicles, although the early Golgi could use Sec22b and GS27/GOSR2 (Hay *et al*., 1997; Lowe *et al*., 1997; Zhang *et al*., 1999; Tai *et al*., 2004), while some TGN and endosome-derived CCD carriers may utilize VAMP4 and STX6 (Bock, Lin and Scheller, 1996; Bock *et al*., 1997; Steegmaier *et al*., 1999; Zeng *et al*., 2003; Tai *et al*., 2004). We also recently discovered the involvement of SNAP29 and VAMP7 in intra-Golgi trafficking (ZD, VL, submitted). Although different classes of CCD carriers are initially recognized at the acceptor cisternae by a specific combination of t-tethers and Rabs (Munro, 2011), the final docking and SNARE pairing of all intra-Golgi vesicles is uniquely orchestrated by the COG complex. COG complex extended 8 tentacle structure allows multiple sequential and/or simultaneous interactions with different components of CCD vesicle tethering and fusion machinery thus stabilizing and proofreading correct pre-fusion arrangements. During acute COG depletion, this final proofreading stage does not occur, causing a massive accumulation of all classes of partially tethered and non-tethered CCD vesicles. The exact mechanism of COG action awaits its resolution, but the use of the novel COG4 acute depletion cellular system unarguably proved that COG machinery is an essential centerpiece of intra-Golgi retrograde trafficking.

**Figure 10:**
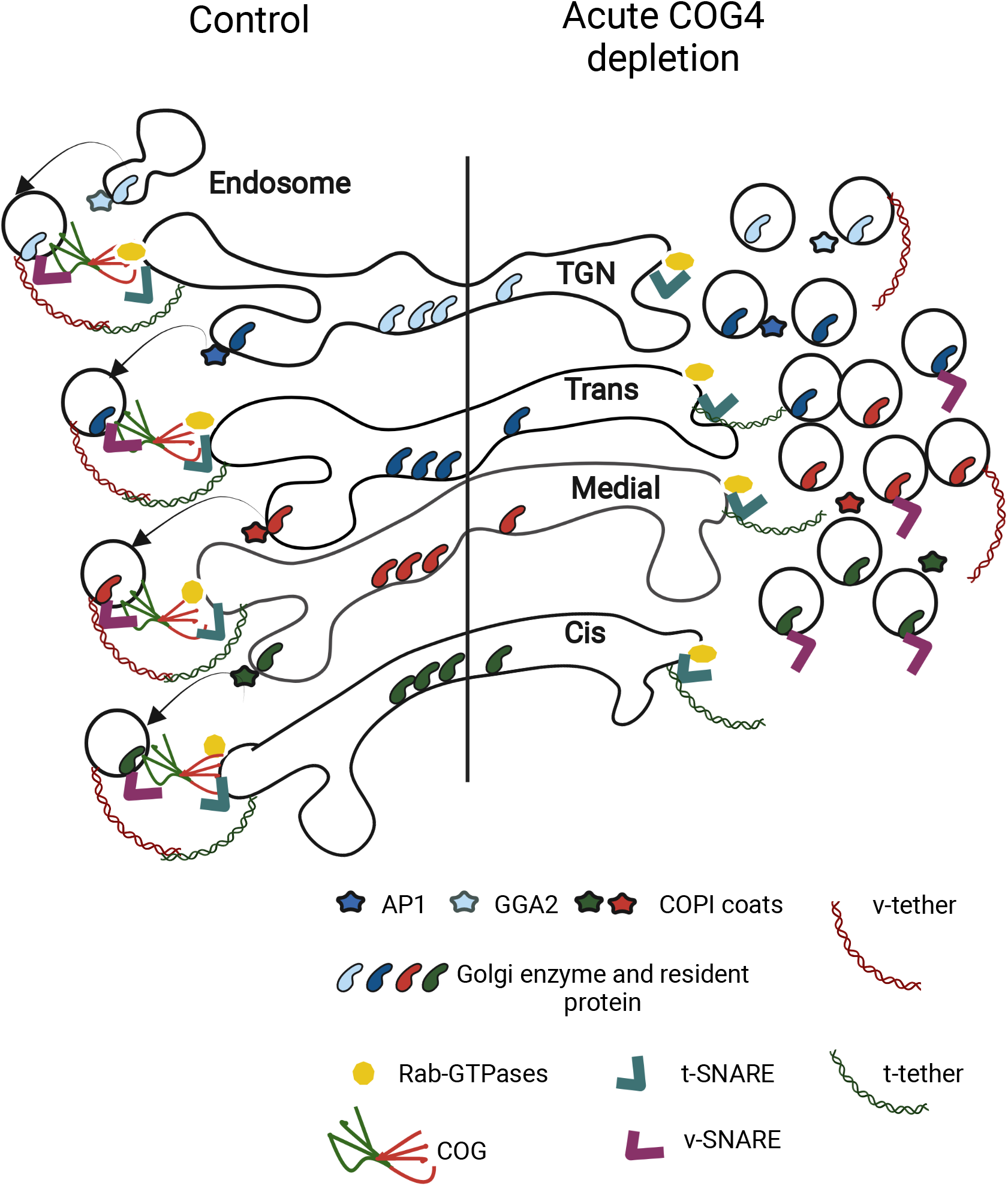
The model depicts the effect of acute COG4 depletion on Golgi homeostasis and vesicular trafficking. Note that upon COG4 depleting t-SNAREs, t-Tethers and Rab proteins remained at the Golgi while vesicular coat protein, glycosylation enzymes, and other Golgi resident proteins dissociated from the Golgi in non-tethered trafficking intermediates.

## Supplementary material

**Supplementary Figure 1.** Rapid COG4 depletion does not cause Golgi fragmentation. Acute COG4 degradation does not displace GM130 from Golgi and does not fragment GM130-positive Golgi indicating normal Golgi morphology. The Airyscan superresolution image of the COG4-mAID clone stained with GM130 (control and 1 hour IAA treatment) was converted to the binary image using ImageJ. The GM130-positive particles having surface area <1um^2 were counted as fragments of Golgi, from n=10 independent experiments with >40 cells were analyzed. Statistical significance was calculated by GraphPad Prism 8 using paired t-test. Here, p ≥ 0.05, non-significant (ns). Error bar represents mean ± SD.

**Supplementary Figure 2.** Acute COG4 depletion differentially alters the abundance of other COG subunits. **(A)** Western blot of time-dependent depletion of COG4-mAID shows the expression of other COG subunits (COG1, COG2, COG3, COG8). 10 µg of total cell lysates were loaded and probed with indicated antibodies. β actin was used as a loading control. **(B)** The graph represents the quantification of **A**. In the bar graph, values represent the mean ± SD from three independent experiments. Statistical significance was calculated using one-way ANOVA. ****p ≤ 0.0001, ***p ≤ 0.001, *p ≤ 0.05, Significant and p ≥ 0.05, non-significant (ns).

**Supplementary Figure 3.** The rapid COG4 depletion differentially alters the total cellular level of Golgi SNAREs. **(A)** Western blot of time-dependent depletion of COG4-mAID shows the expression of Golgi SNAREs (GS15, GS28, STX5, YKT6). 10 µg of total cell lysates were loaded and probed with indicated antibodies. Β actin was used as a loading control. **(B)** The graph represents the quantification of **A**. In the bar graph, values represent the mean ± SD from three independent experiments. Statistical significance was calculated using one-way ANOVA. P ≥ 0.05, non-significant (ns), ****p ≤ 0.0001, **p ≤ 0.01, significant.

**Supplementary Figure 4.** The rapid COG4 depletion partially displaces TMF1 from Golgi but no effect on localization of trans-Golgi golgin97 and golgin245 respectively. Airyscan superresolution IF analysis of untreated (control) or IAA treated COG4-mAID cells stained for **(A)** GM130 (green) and TMF1 (red), **(B)** GM130 (green) and Golgin97 (red), and **(C)** GM130 (green) and golgin245 (red). Scale bars, 20 µm. For the better presentation green and red channels are shown in inverted black and white mode whereas the merged view is shown in RGB mode. **(D)** Colocalization of tested golgins with GM130 was determined by using Pearson’s correlation coefficient, >40 cells were analyzed. Statistical significance was calculated by GraphPad Prism 8 using paired t-test. Here, p ≥ 0.05, non-significant (ns), ****p ≤ 0.0001 (significant). Error bar represents mean ± SD.

**Supplementary Figure 5.** Acute COG4 depletion does not alter the localization of exogenously overexpressed GFP-Rab30A and GFP-Rab43B. Airyscan superresolution IF analysis of untreated (control) or IAA treated COG4-mAID cells transiently transfected with corresponding GFP-tagged Rabs and stained for **(A)** GM130 (red) and GFP-Rab30A, **(B)** GM130 (red), and GFP-Rab43A. Scale bars, 10 µm. For better presentation green and red channels are shown in inverted black and white mode whereas the merged view is shown in RGB mode. **(C)** Colocalization of GM130 and GFP-RabGTPases was performed using Pearson’s correlation coefficient, >20 cells were analyzed. Here, p ≥ 0.05, nonsignificant (ns). Error bar represents mean ± SD.

**Supplementary Figure 6**. B4GalT1 and GPP130 recycle in different CCD vesicles **(A)** Airyscan superresolution IF analysis of untreated (control) or IAA treated COG4-mAID cells stained for B4GalT1 (red) and GPP130 (green). Scale bars, 20 µm. The enlarged (5x) view of the framed area of the upper panel is shown in the bottom panel. For better presentation green and red channels are shown in inverted black and white mode whereas the merged view is shown in RGB mode. **(B)** Colocalization of B4GalT1 with GPP130 was determined by calculating Pearson’s correlation coefficient, >90 cells were analyzed. Statistical significance was calculated by GraphPad Prism 8 using paired t-test. Here, *p ≤ 0.05, (significant). Error bar represents mean ± SD. **(C)** Western blot of sucrose gradient fractions shows the expression of B4GalT1 and GPP130 by probing with corresponding antibodies. **(D)** The line graph represents the quantification of **C.**

**Supplementary Figure 7.** The acute COG4 knockdown is displacing cargo adaptor GOLPH3 from Golgi**. (A)** Airyscan superresolution IF analysis of untreated (control) or IAA treated COG4-mAID cells stained for GM130 (green) and GOLPH3 (red). Scale bars, 20 µm. For better presentation green and red channels are shown in inverted black and white mode whereas the merged view is shown in RGB mode. **(B)** Colocalization of GOLPH3 with GM130 was determined by calculating Pearson’s correlation coefficient, >90 cells were analyzed. Statistical significance was calculated by GraphPad Prism 8 using paired t-test. Here, ****p ≤ 0.0001 (significant). Error bar represents mean ± SD.

## Author Contributions

F.T.S wrote the article and made substantial contributions to conception and design, acquisition of data, analysis, and interpretation of data. I.D.P. participated in drafting the article, performed EM and DNA cloning experiments, and interpreted the data. Z.D. cloned COG4 promoter region and edited the article. V.L. edited the article and made substantial contributions to conception and design.

## Acknowledgments

We are thankful to Rainer Duden, Elizabeth Sztul, Graham Warren as well as others who provided reagents. We are thankful to Tetyana Kudlyk for excellent technical support and Amrita Khakurel for critical discussion. We would also like to thank the UAMS Digital Microscopy, sequencing and flow cytometry core facilities for the use of their facilities and expertise.

## References

Adolf, F. et al. (2019) ‘Proteomic Profiling of Mammalian COPII and COPI Vesicles’, Cell Reports, 26(1), pp. 250–265.e5. doi:10.1016/j.celrep.2018.12.041.

Bailey Blackburn, J., et al. (2016) ‘COG Complex Complexities: Detailed Characterization of a Complete Set of HEK293T Cells Lacking Individual COG Subunits’, Frontiers in Cell and Developmental Biology, 4. doi:10.3389/fcell.2016.00023.

Bentley, M. et al. (2006) ‘SNARE status regulates tether recruitment and function in homotypic COPII vesicle fusion’, The Journal of Biological Chemistry, 281(50), pp. 38825–38833. doi:10.1074/jbc.M606044200.

Berninsone, P.M. and Hirschberg, C.B. (2000) ‘Nucleotide sugar transporters of the Golgi apparatus’, Current Opinion in Structural Biology, 10(5), pp. 542–547. doi:10.1016/s0959-440x(00)00128-7.

Béthune, J. et al. (2006) ‘Coatomer, the coat protein of COPI transport vesicles, discriminates endoplasmic reticulum residents from p24 proteins’, Molecular and Cellular Biology, 26(21), pp. 8011–8021. doi:10.1128/MCB.01055-06.

Beznoussenko, G.V. et al. (2014) ‘Transport of soluble proteins through the Golgi occurs by diffusion via continuities across cisternae’, eLife, 3. doi:10.7554/eLife.02009.

Bhuin, T. and Roy, J.K. (2014) ‘Rab proteins: the key regulators of intracellular vesicle transport’, Experimental Cell Research, 328(1), pp. 1–19. doi:10.1016/j.yexcr.2014.07.027.

Blackburn, J.B. et al. (2018) ‘More than just sugars: COG complex deficiency causes glycosylation-independent cellular defects’, Traffic (Copenhagen, Denmark), 19(6), pp. 463– 480. doi:10.1111/tra.12564.

Blackburn, J.B., D’Souza, Z. and Lupashin, V.V. (2019) ‘Maintaining order: COG complex controls Golgi trafficking, processing, and sorting’, FEBS Letters, 593(17), pp. 2466–2487. doi:10.1002/1873-3468.13570.

Blackburn, J.B. and Lupashin, V.V. (2016) ‘Creating Knockouts of Conserved Oligomeric Golgi Complex Subunits Using CRISPR-Mediated Gene Editing Paired with a Selection Strategy Based on Glycosylation Defects Associated with Impaired COG Complex Function’, Methods in Molecular Biology (Clifton, N.J.), 1496, pp. 145–161. doi:10.1007/978-1-4939-6463-5_12.

Bock, J.B. et al. (1997) ‘Syntaxin 6 functions in trans-Golgi network vesicle trafficking’, Molecular Biology of the Cell, 8(7), pp. 1261–1271. doi:10.1091/mbc.8.7.1261.

Bock, J.B., Lin, R.C. and Scheller, R.H. (1996) ‘A new syntaxin family member implicated in targeting of intracellular transport vesicles’, The Journal of Biological Chemistry, 271(30), pp. 17961–17965. doi:10.1074/jbc.271.30.17961.

Boncompain, G. et al. (2012) ‘Synchronization of secretory protein traffic in populations of cells’, Nature Methods, 9(5), pp. 493–498. doi:10.1038/nmeth.1928.

Brooks, S.A. (2000) ‘The involvement of Helix pomatia lectin (HPA) binding N-acetylgalactosamine glycans in cancer progression’, Histology and Histopathology, 15(1), pp. 143–158. doi:10.14670/HH-15.143.

Cai, H., Reinisch, K. and Ferro-Novick, S. (2007) ‘Coats, tethers, Rabs, and SNAREs work together to mediate the intracellular destination of a transport vesicle’, Developmental Cell, 12(5), pp. 671–682. doi:10.1016/j.devcel.2007.04.005.

Cao, X. and Barlowe, C. (2000) ‘Asymmetric requirements for a Rab GTPase and SNARE proteins in fusion of COPII vesicles with acceptor membranes’, The Journal of Cell Biology, 149(1), pp. 55–66. doi:10.1083/jcb.149.1.55.

Casler, J.C. et al. (2019) ‘Maturation-driven transport and AP-1-dependent recycling of a secretory cargo in the Golgi’, The Journal of Cell Biology, 218(5), pp. 1582–1601. doi:10.1083/jcb.201807195.

Climer, L.K. et al. (2018) ‘Membrane detachment is not essential for COG complex function’, Molecular Biology of the Cell, 29(8), pp. 964–974. doi:10.1091/mbc.E17-11-0694.

Costaguta, G. et al. (2001) ‘Yeast Gga coat proteins function with clathrin in Golgi to endosome transport’, Molecular Biology of the Cell, 12(6), pp. 1885–1896. doi:10.1091/mbc.12.6.1885.

Cottam, N.P. and Ungar, D. (2012) ‘Retrograde vesicle transport in the Golgi’, Protoplasma, 249(4), pp. 943–955. doi:10.1007/s00709-011-0361-7.

Daboussi, L. et al. (2017) ‘Conserved role for Gga proteins in phosphatidylinositol 4-kinase localization to the trans-Golgi network’, Proceedings of the National Academy of Sciences of the United States of America, 114(13), pp. 3433–3438. doi:10.1073/pnas.1615163114.

Dan, X., Liu, W. and Ng, T.B. (2016) ‘Development and Applications of Lectins as Biological Tools in Biomedical Research’, Medicinal Research Reviews, 36(2), pp. 221–247. doi:10.1002/med.21363.

Dharmasiri, N. and Estelle, M. (2004) ‘Auxin signaling and regulated protein degradation’, Trends in Plant Science, 9(6), pp. 302–308. doi:10.1016/j.tplants.2004.04.003.

Di Martino, R., Sticco, L. and Luini, A. (2019) ‘Regulation of cargo export and sorting at the trans-Golgi network’, FEBS letters, 593(17), pp. 2306–2318. doi:10.1002/1873-3468.13572.

D’Souza, Z. et al. (2019) ‘Defects in COG-Mediated Golgi Trafficking Alter Endo-Lysosomal System in Human Cells’, Frontiers in Cell and Developmental Biology, 7, p. 118. doi:10.3389/fcell.2019.00118.

D’Souza, Z. et al. (2021) ‘Getting Sugar Coating Right! The Role of the Golgi Trafficking Machinery in Glycosylation’, Cells, 10(12), p. 3275. doi:10.3390/cells10123275.

D’Souza, Z., Taher, F.S. and Lupashin, V.V. (2020) ‘Golgi inCOGnito: From vesicle tethering to human disease’, Biochimica et Biophysica Acta (BBA) - General Subjects, 1864(11), p. 129694. doi:10.1016/j.bbagen.2020.129694.

Dull, T. et al. (1998) ‘A third-generation lentivirus vector with a conditional packaging system’, Journal of Virology, 72(11), pp. 8463–8471. doi:10.1128/JVI.72.11.8463-8471.1998.

Duncan, M.C. (2022) ‘New directions for the clathrin adaptor AP-1 in cell biology and human disease’, Current Opinion in Cell Biology, 76, p. 102079. doi:10.1016/j.ceb.2022.102079.

Fotso, P. et al. (2005) ‘Cog1p Plays a Central Role in the Organization of the Yeast Conserved Oligomeric Golgi Complex *’, Journal of Biological Chemistry, 280(30), pp. 27613–27623. doi:10.1074/jbc.M504597200.

Foulquier, F. (2009) ‘COG defects, birth and rise!’, Biochimica et Biophysica Acta (BBA) - Molecular Basis of Disease, 1792(9), pp. 896–902. doi:10.1016/j.bbadis.2008.10.020.

Fridmann-Sirkis, Y., Siniossoglou, S. and Pelham, H.R.B. (2004) ‘TMF is a golgin that binds Rab6 and influences Golgi morphology’, BMC cell biology, 5, p. 18. doi:10.1186/1471-2121-5-18.

Gilchrist, A. et al. (2006) ‘Quantitative proteomics analysis of the secretory pathway’, Cell, 127(6), pp. 1265–1281. doi:10.1016/j.cell.2006.10.036.

Glick, B.S. and Nakano, A. (2009) ‘Membrane traffic within the Golgi apparatus’, Annual Review of Cell and Developmental Biology, 25, pp. 113–132. doi:10.1146/annurev.cellbio.24.110707.175421.

Gu, F., Crump, C.M. and Thomas, G. (2001) ‘Trans-Golgi network sorting’, Cellular and molecular life sciences: CMLS, 58(8), pp. 1067–1084. doi:10.1007/PL00000922.

Hay, J.C. et al. (1997) ‘Protein interactions regulating vesicle transport between the endoplasmic reticulum and Golgi apparatus in mammalian cells’, Cell, 89(1), pp. 149–158. doi:10.1016/s0092-8674(00)80191-9.

Holland, A.J. et al. (2012) ‘Inducible, reversible system for the rapid and complete degradation of proteins in mammalian cells’, Proceedings of the National Academy of Sciences of the United States of America, 109(49), pp. E3350–E3357. doi:10.1073/pnas.1216880109.

Huang, S. and Wang, Y. (2017) ‘Golgi structure formation, function, and post-translational modifications in mammalian cells’, F1000Research, 6, p. 2050. doi:10.12688/f1000research.11900.1.

Hwang, I. (2008) ‘Sorting and anterograde trafficking at the Golgi apparatus’, Plant Physiology, 148(2), pp. 673–683. doi:10.1104/pp.108.124925.

Jaquinod, M. et al. (2007) ‘A proteomics dissection of Arabidopsis thaliana vacuoles isolated from cell culture’, Molecular & cellular proteomics: MCP, 6(3), pp. 394–412. doi:10.1074/mcp.M600250-MCP200.

Jin, H. et al. (2021) ‘Rab GTPases: Central Coordinators of Membrane Trafficking in Cancer’, Frontiers in Cell and Developmental Biology, 9, p. 648384. doi:10.3389/fcell.2021.648384.

Kaku, H. and Goldstein, I.J. (1992) ‘Interaction of linear manno-oligosaccharides with three mannose-specific bulb lectins. Comparison with mannose/glucose-binding lectins’, Carbohydrate Research, 229(2), pp. 337–346. doi:10.1016/s0008-6215(00)90579-2.

Khakurel, A. et al. (2021) ‘The Golgi-associated retrograde protein (GARP) complex plays an essential role in the maintenance of the Golgi glycosylation machinery’, Molecular Biology of the Cell. Edited by B.S. Glick, p. mbc21-04-0169. doi:10.1091/mbc.E21-04-0169.

Kim, D.W. et al. (2001) ‘Sgf1p, a new component of the Sec34p/Sec35p complex’, Traffic (Copenhagen, Denmark), 2(11), pp. 820–830. doi:10.1034/j.1600-0854.2001.21111.x.

Kingsley, D.M. et al. (1986) ‘Three types of low density lipoprotein receptor-deficient mutant have pleiotropic defects in the synthesis of N-linked, O-linked, and lipid-linked carbohydrate chains’, The Journal of Cell Biology, 102(5), pp. 1576–1585. doi:10.1083/jcb.102.5.1576.

Kudlyk, T. et al. (2013) ‘COG6 interacts with a subset of the Golgi SNAREs and is important for the Golgi complex integrity’, Traffic (Copenhagen, Denmark), 14(2), pp. 194–204. doi:10.1111/tra.12020.

Kweon, H.-S. et al. (2004) ‘Golgi enzymes are enriched in perforated zones of golgi cisternae but are depleted in COPI vesicles’, Molecular Biology of the Cell, 15(10), pp. 4710–4724. doi:10.1091/mbc.e03-12-0881.

Laufman, O. et al. (2009) ‘Direct interaction between the COG complex and the SM protein, Sly1, is required for Golgi SNARE pairing’, The EMBO journal, 28(14), pp. 2006–2017. doi:10.1038/emboj.2009.168.

Laufman, O., Hong, W. and Lev, S. (2013) ‘The COG complex interacts with multiple Golgi SNAREs and enhances fusogenic assembly of SNARE complexes’, Journal of Cell Science, 126(6), pp. 1506–1516. doi:10.1242/jcs.122101.

Lees, J.A. et al. (2010) ‘Molecular organization of the COG vesicle tethering complex’, Nature structural & molecular biology, 17(11), pp. 1292–1297. doi:10.1038/nsmb.1917.

Li, G. and Marlin, M.C. (2015) ‘Rab family of GTPases’, Methods in Molecular Biology (Clifton, N.J.), 1298, pp. 1–15. doi:10.1007/978-1-4939-2569-8_1.

Linders, P.T. et al. (2019) ‘Stx5-Mediated ER-Golgi Transport in Mammals and Yeast’, Cells, 8(8), p. E780. doi:10.3390/cells8080780.

Linders, P.T.A. et al. (2020) ‘Sugary Logistics Gone Wrong: Membrane Trafficking and Congenital Disorders of Glycosylation’, International Journal of Molecular Sciences, 21(13), p. 4654. doi:10.3390/ijms21134654.

Liu, L., Doray, B. and Kornfeld, S. (2018) ‘Recycling of Golgi glycosyltransferases requires direct binding to coatomer’, Proceedings of the National Academy of Sciences of the United States of America, 115(36), pp. 8984–8989. doi:10.1073/pnas.1810291115.

Liu, S. and Storrie, B. (2012) ‘Are Rab proteins the link between Golgi organization and membrane trafficking?’, Cellular and molecular life sciences: CMLS, 69(24), pp. 4093–4106. doi:10.1007/s00018-012-1021-6.

Love, H.D. et al. (1998) ‘Isolation of functional Golgi-derived vesicles with a possible role in retrograde transport’, The Journal of Cell Biology, 140(3), pp. 541–551. doi:10.1083/jcb.140.3.541.

Lowe, S.L. et al. (1997) ‘A SNARE involved in protein transport through the Golgi apparatus’, Nature, 389(6653), pp. 881–884. doi:10.1038/39923.

Luo, P.M. and Boyce, M. (2019) ‘Directing Traffic: Regulation of COPI Transport by Post-translational Modifications’, Frontiers in Cell and Developmental Biology, 7, p. 190. doi:10.3389/fcell.2019.00190.

Malsam, J. et al. (2005) ‘Golgin tethers define subpopulations of COPI vesicles’, Science (New York, N.Y.), 307(5712), pp. 1095–1098. doi:10.1126/science.1108061.

Miller, V.J. et al. (2013) ‘Molecular Insights into Vesicle Tethering at the Golgi by the Conserved Oligomeric Golgi (COG) Complex and the Golgin TATA Element Modulatory Factor (TMF)’, The Journal of Biological Chemistry, 288(6), pp. 4229–4240. doi:10.1074/jbc.M112.426767.

Mironov, A.A. and Beznoussenko, G.V. (2019) ‘Models of Intracellular Transport: Pros and Cons’, Frontiers in Cell and Developmental Biology, 7, p. 146. doi:10.3389/fcell.2019.00146.

Munro, S. (2011) ‘The golgin coiled-coil proteins of the Golgi apparatus’, Cold Spring Harbor Perspectives in Biology, 3(6), p. a005256. doi:10.1101/cshperspect.a005256.

Natsume, T. et al. (2016) ‘Rapid Protein Depletion in Human Cells by Auxin-Inducible Degron Tagging with Short Homology Donors’, Cell Reports, 15(1), pp. 210–218. doi:10.1016/j.celrep.2016.03.001.

Nilsson, T. et al. (1993) ‘Overlapping distribution of two glycosyltransferases in the Golgi apparatus of HeLa cells’, The Journal of Cell Biology, 120(1), pp. 5–13. doi:10.1083/jcb.120.1.5.

Nishimura, K. et al. (2009) ‘An auxin-based degron system for the rapid depletion of proteins in nonplant cells’, Nature Methods, 6(12), pp. 917–922. doi:10.1038/nmeth.1401.

Oka, T. et al. (2004) ‘The COG and COPI complexes interact to control the abundance of GEARs, a subset of Golgi integral membrane proteins’, Molecular Biology of the Cell, 15(5), pp. 2423–2435. doi:10.1091/mbc.e03-09-0699.

Oka, T. et al. (2005) ‘Genetic Analysis of the Subunit Organization and Function of the Conserved Oligomeric Golgi (COG) Complex: STUDIES OF COG5- AND COG7-DEFICIENT MAMMALIAN CELLS*’, Journal of Biological Chemistry, 280(38), pp. 32736–32745. doi:10.1074/jbc.M505558200.

Ondruskova, N. et al. (2021) ‘Congenital disorders of glycosylation: Still “hot” in 2020’, Biochimica Et Biophysica Acta. General Subjects, 1865(1), p. 129751. doi:10.1016/j.bbagen.2020.129751.

Orci, L. et al. (1997) ‘Bidirectional transport by distinct populations of COPI-coated vesicles’, Cell, 90(2), pp. 335–349. doi:10.1016/s0092-8674(00)80341-4.

Orci, L., Ravazzola, M., et al. (2000) ‘Anterograde flow of cargo across the golgi stack potentially mediated via bidirectional “percolating” COPI vesicles’, Proceedings of the National Academy of Sciences of the United States of America, 97(19), pp. 10400–10405. doi:10.1073/pnas.190292497.

Orci, L., Amherdt, M., et al. (2000) ‘Exclusion of golgi residents from transport vesicles budding from Golgi cisternae in intact cells’, The Journal of Cell Biology, 150(6), pp. 1263–1270. doi:10.1083/jcb.150.6.1263.

Park, K. et al. (2021) ‘The Golgi complex: a hub of the secretory pathway’, BMB reports, 54(5), pp. 246–252.

Peanne, R. et al. (2011) ‘Differential effects of lobe A and lobe B of the Conserved Oligomeric Golgi complex on the stability of β1,4-galactosyltransferase 1 and α2,6-sialyltransferase 1’, Glycobiology, 21(7), pp. 864–876. doi:10.1093/glycob/cwq176.

Pelham, H.R. (2001) ‘Traffic through the Golgi apparatus’, The Journal of Cell Biology, 155(7), pp. 1099–1101. doi:10.1083/jcb.200110160.

Petitjean, O. et al. (2020) ‘Genome-Wide CRISPR-Cas9 Screen Reveals the Importance of the Heparan Sulfate Pathway and the Conserved Oligomeric Golgi Complex for Synthetic Double-Stranded RNA Uptake and Sindbis Virus Infection’, mSphere, 5(6), pp. e00914–20. doi:10.1128/mSphere.00914-20.

Pokrovskaya, Irina et al. (2011) ‘COG Complex Specifically Regulates the Maintenance of Golgi Glycosylation Machinery’, Glycobiology, 21, pp. 1554–69. doi:10.1093/glycob/cwr028.

Pokrovskaya, I. et al. (2011) ‘Conserved oligomeric Golgi complex specifically regulates the maintenance of Golgi glycosylation machinery.’, Glycobiology [Preprint]. doi:10.1093/glycob/cwr028.

Popoff, V. et al. (2011) ‘COPI budding within the Golgi stack’, Cold Spring Harbor Perspectives in Biology, 3(11), p. a005231. doi:10.1101/cshperspect.a005231.

Rabouille, C. et al. (1995) ‘Mapping the distribution of Golgi enzymes involved in the construction of complex oligosaccharides’, Journal of Cell Science, 108 ( Pt 4), pp. 1617–1627. doi:10.1242/jcs.108.4.1617.

Ram, R.J., Li, B. and Kaiser, C.A. (2002) ‘Identification of Sec36p, Sec37p, and Sec38p: components of yeast complex that contains Sec34p and Sec35p’, Molecular Biology of the Cell, 13(5), pp. 1484–1500. doi:10.1091/mbc.01-10-0495.

Reynders, E. et al. (2011) ‘How Golgi glycosylation meets and needs trafficking: the case of the COG complex’, Glycobiology, 21(7), pp. 853–863. doi:10.1093/glycob/cwq179.

Richardson, B.C. et al. (2009) ‘Structural basis for a human glycosylation disorder caused by mutation of the COG4 gene’, Proceedings of the National Academy of Sciences, 106(32), pp. 13329–13334. doi:10.1073/pnas.0901966106.

Rizzo, R. et al. (2013) ‘The dynamics of engineered resident proteins in the mammalian Golgi complex relies on cisternal maturation’, The Journal of Cell Biology, 201(7), pp. 1027–1036. doi:10.1083/jcb.201211147.

Robinson, M.S. (2004) ‘Adaptable adaptors for coated vesicles’, Trends in Cell Biology, 14(4), pp. 167–174. doi:10.1016/j.tcb.2004.02.002.

Röttger, S. et al. (1998) ‘Localization of three human polypeptide GalNAc-transferases in HeLa cells suggests initiation of O-linked glycosylation throughout the Golgi apparatus’, Journal of Cell Science, 111 ( Pt 1), pp. 45–60. doi:10.1242/jcs.111.1.45.

Rowe, T. et al. (1998) ‘Role of vesicle-associated syntaxin 5 in the assembly of pre-Golgi intermediates’, Science (New York, N.Y.), 279(5351), pp. 696–700. doi:10.1126/science.279.5351.696.

Schmitz, K.R. et al. (2008) ‘Golgi Localization of Glycosyltransferases Requires a Vps74p Oligomer’, Developmental Cell, 14(4), pp. 523–534. doi:10.1016/j.devcel.2008.02.016.

Shestakova, A. et al. (2007) ‘Interaction of the conserved oligomeric Golgi complex with t-SNARE Syntaxin5a/Sed5 enhances intra-Golgi SNARE complex stability’, The Journal of Cell Biology, 179(6), pp. 1179–1192. doi:10.1083/jcb.200705145.

Shestakova, A., Zolov, S. and Lupashin, V. (2006) ‘COG complex-mediated recycling of Golgi glycosyltransferases is essential for normal protein glycosylation’, Traffic (Copenhagen, Denmark), 7(2), pp. 191–204. doi:10.1111/j.1600-0854.2005.00376.x.

Smith, R.D. et al. (2009) ‘The COG Complex, Rab6 and COPI Define a Novel Golgi Retrograde Trafficking Pathway that is Exploited by SubAB Toxin’, Traffic, 10(10), pp. 1502–1517. doi:10.1111/j.1600-0854.2009.00965.x.

Sohda, M. et al. (2007) ‘The Interaction of Two Tethering Factors, p115 and COG complex, is Required for Golgi Integrity’, Traffic, 8(3), pp. 270–284. doi:10.1111/j.1600-0854.2006.00530.x.

Sohda, M. et al. (2010) ‘Interaction of Golgin-84 with the COG Complex Mediates the Intra-Golgi Retrograde Transport’, Traffic, 11(12), pp. 1552–1566. doi:10.1111/j.1600-0854.2010.01123.x.

Stanley, P. (2011) ‘Golgi glycosylation’, Cold Spring Harbor Perspectives in Biology, 3(4), p. a005199. doi:10.1101/cshperspect.a005199.

Steegmaier, M. et al. (1999) ‘Vesicle-associated membrane protein 4 is implicated in trans-Golgi network vesicle trafficking’, Molecular Biology of the Cell, 10(6), pp. 1957–1972. doi:10.1091/mbc.10.6.1957.

Steet, R. and Kornfeld, S. (2006) ‘COG-7-deficient Human Fibroblasts Exhibit Altered Recycling of Golgi Proteins’, Molecular Biology of the Cell, 17(5), pp. 2312–2321. doi:10.1091/mbc.e05-08-0822.

Stewart, S.A. et al. (2003) ‘Lentivirus-delivered stable gene silencing by RNAi in primary cells’, RNA, 9(4), pp. 493–501. doi:10.1261/rna.2192803.

Sumya, F.T., Pokrovskaya, I.D. and Lupashin, V. (2021) ‘Development and Initial Characterization of Cellular Models for COG Complex-Related CDG-II Diseases’, Frontiers in Genetics, 12, p. 733048. doi:10.3389/fgene.2021.733048.

Suvorova, E.S., Duden, R. and Lupashin, V.V. (2002) ‘The Sec34/Sec35p complex, a Ypt1p effector required for retrograde intra-Golgi trafficking, interacts with Golgi SNAREs and COPI vesicle coat proteins’, Journal of Cell Biology, 157(4), pp. 631–643. doi:10.1083/jcb.200111081.

Tai, G. et al. (2004) ‘Participation of the syntaxin 5/Ykt6/GS28/GS15 SNARE complex in transport from the early/recycling endosome to the trans-Golgi network’, Molecular Biology of the Cell, 15(9), pp. 4011–4022. doi:10.1091/mbc.e03-12-0876.

Tu, L. et al. (2008) ‘Signal-mediated dynamic retention of glycosyltransferases in the Golgi’, Science (New York, N.Y.), 321(5887), pp. 404–407. doi:10.1126/science.1159411.

Ungar, D. et al. (2002) ‘Characterization of a mammalian Golgi-localized protein complex, COG, that is required for normal Golgi morphology and function’, The Journal of Cell Biology, 157(3), pp. 405–415. doi:10.1083/jcb.200202016.

Ungar, D. et al. (2005) ‘Subunit architecture of the conserved oligomeric Golgi complex’, The Journal of Biological Chemistry, 280(38), pp. 32729–32735. doi:10.1074/jbc.M504590200.

Ungar, D. et al. (2006) ‘Retrograde transport on the COG railway’, Trends in Cell Biology, 16(2), pp. 113–120. doi:10.1016/j.tcb.2005.12.004.

Welch, L.G. et al. (2021) ‘GOLPH3 and GOLPH3L are broad-spectrum COPI adaptors for sorting into intra-Golgi transport vesicles’, Journal of Cell Biology, 220(10), p. e202106115. doi:10.1083/jcb.202106115.

Willett, R. et al. (2013) ‘COG complexes form spatial landmarks for distinct SNARE complexes’, Nature Communications, 4, p. 1553. doi:10.1038/ncomms2535.

Willett, R. et al. (2014) ‘Multipronged interaction of the COG complex with intracellular membranes’, Cellular Logistics, 4, p. e27888. doi:10.4161/cl.27888.

Willett, R. et al. (2016) ‘COG lobe B sub-complex engages v-SNARE GS15 and functions via regulated interaction with lobe A sub-complex’, Scientific Reports, 6(1), p. 29139. doi:10.1038/srep29139.

Willett, R., Ungar, D. and Lupashin, V. (2013) ‘The Golgi puppet master: COG complex at center stage of membrane trafficking interactions’, Histochemistry and Cell Biology, 140(3), pp. 271–283. doi:10.1007/s00418-013-1117-6.

Wong, M. and Munro, S. (2014) ‘Membrane trafficking. The specificity of vesicle traffic to the Golgi is encoded in the golgin coiled-coil proteins’, Science (New York, N.Y.), 346(6209), p. 1256898. doi:10.1126/science.1256898.

Zeevaert, R. et al. (2008) ‘Deficiencies in subunits of the Conserved Oligomeric Golgi (COG) complex define a novel group of Congenital Disorders of Glycosylation’, Molecular Genetics and Metabolism, 93(1), pp. 15–21. doi:10.1016/j.ymgme.2007.08.118.

Zeng, Q. et al. (2003) ‘The cytoplasmic domain of Vamp4 and Vamp5 is responsible for their correct subcellular targeting: the N-terminal extenSion of VAMP4 contains a dominant autonomous targeting signal for the trans-Golgi network’, The Journal of Biological Chemistry, 278(25), pp. 23046–23054. doi:10.1074/jbc.M303214200.

Zhang, T. et al. (1999) ‘Morphological and functional association of Sec22b/ERS-24 with the pre-Golgi intermediate compartment’, Molecular Biology of the Cell, 10(2), pp. 435–453. doi:10.1091/mbc.10.2.435.

Zhang, T. and Hong, W. (2001) ‘Ykt6 forms a SNARE complex with syntaxin 5, GS28, and Bet1 and participates in a late stage in endoplasmic reticulum-Golgi transport’, The Journal of Biological Chemistry, 276(29), pp. 27480–27487. doi:10.1074/jbc.M102786200.

Zhang, Y. and Seemann, J. (2021) ‘Rapid degradation of GRASP55 and GRASP65 reveals their immediate impact on the Golgi structure’, Journal of Cell Biology, 220(1), p. e202007052. doi:10.1083/jcb.202007052.

Zolov, S.N. and Lupashin, V.V. (2005) ‘Cog3p depletion blocks vesicle-mediated Golgi retrograde trafficking in HeLa cells’, The Journal of Cell Biology, 168(5), pp. 747–759. doi:10.1083/jcb.200412003.

