## Supplementary figures and images for "Acute COG inactivation unveiled its immediate impact on Golgi and illuminated the nature of intra-Golgi recycling vesicles"

### Supplemental Figure 1

**Supplementary Figure 1**

**A)**

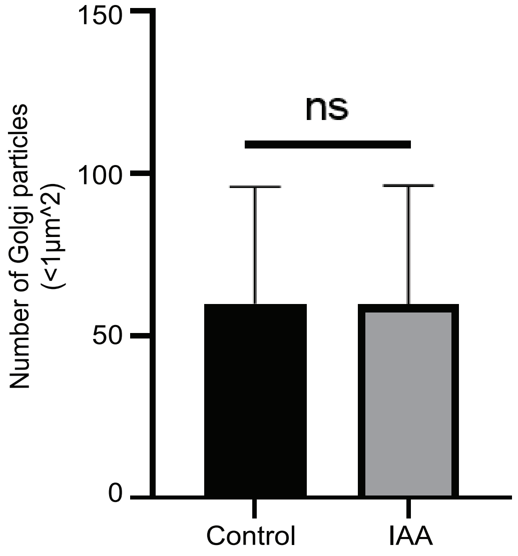

### Supplemental Figure 2

**Supplementary 2**

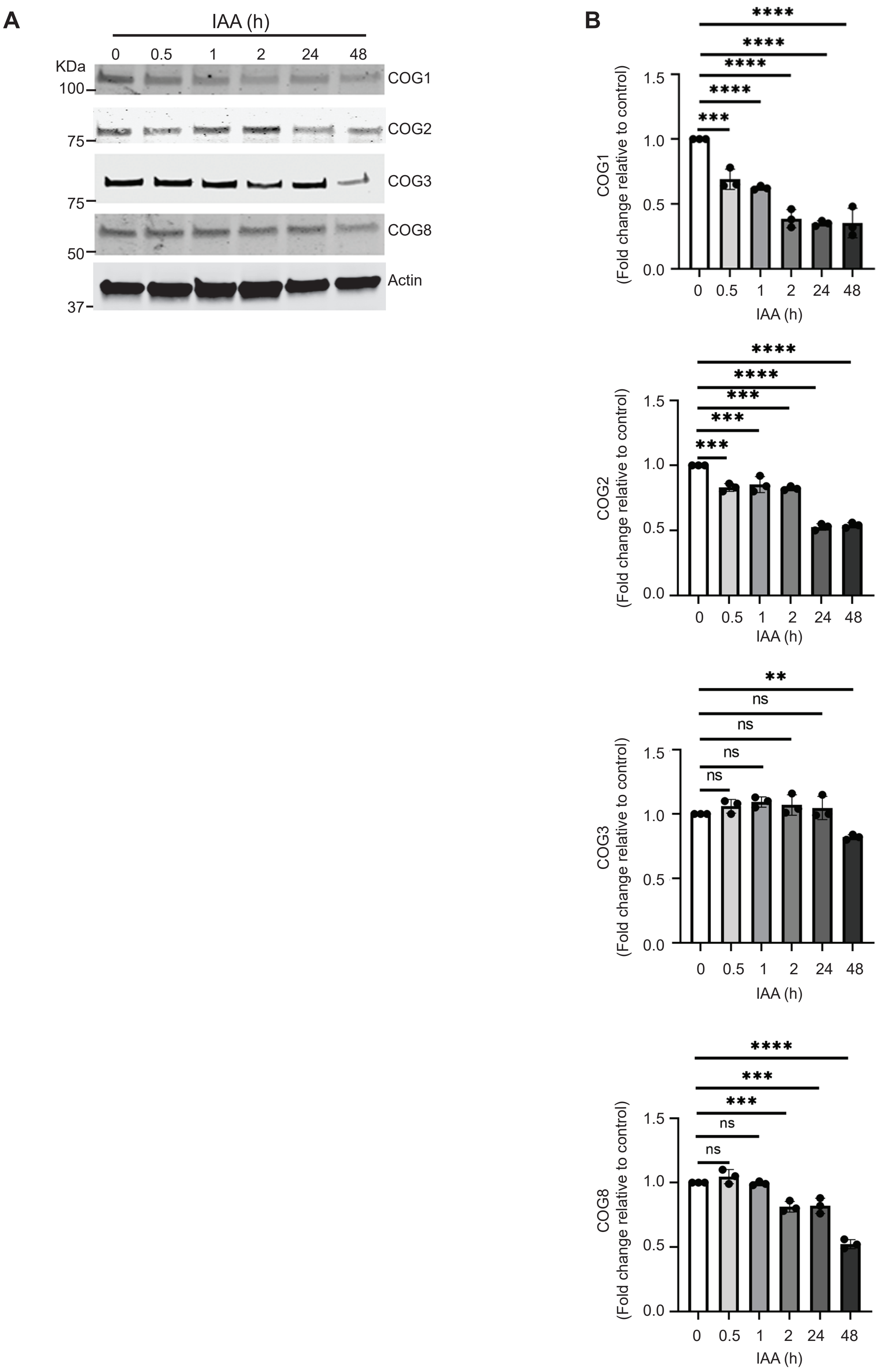

### Supplemental Figure 3

**Supplementary 3**

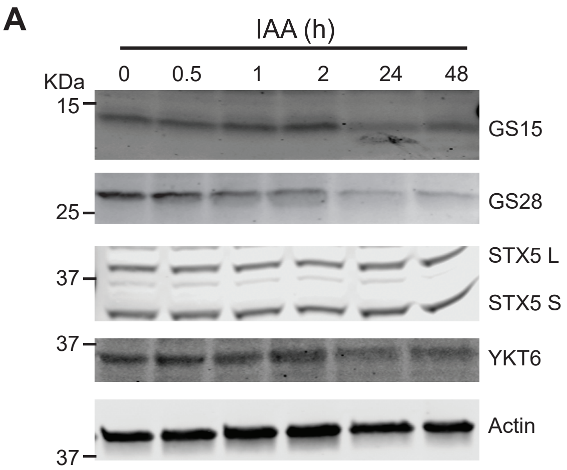


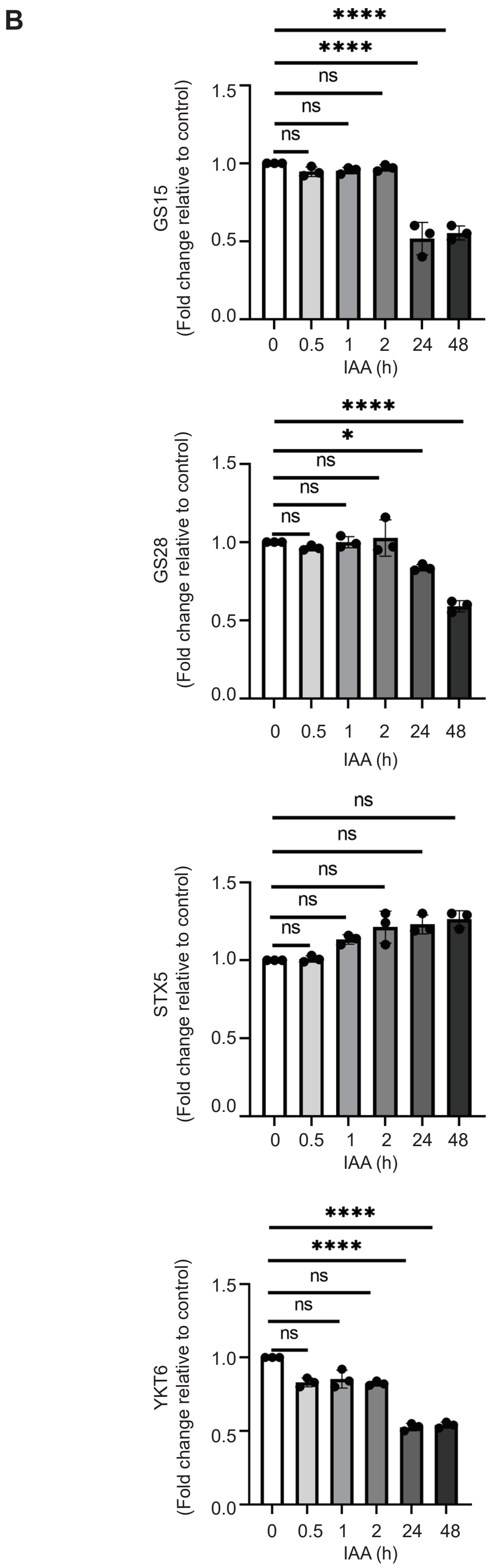

### Supplemental Figure 4

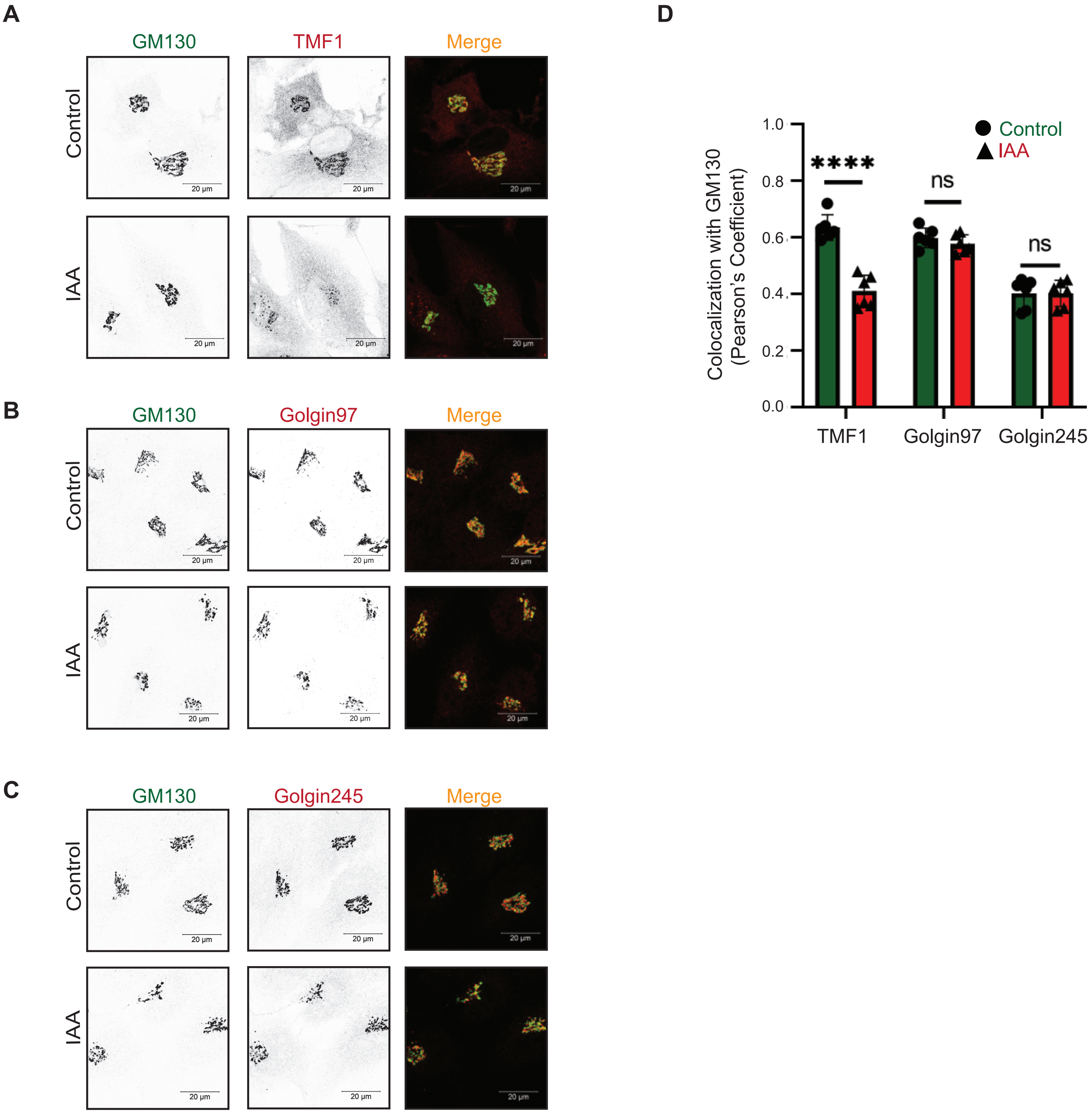

### Supplemental Figure 5

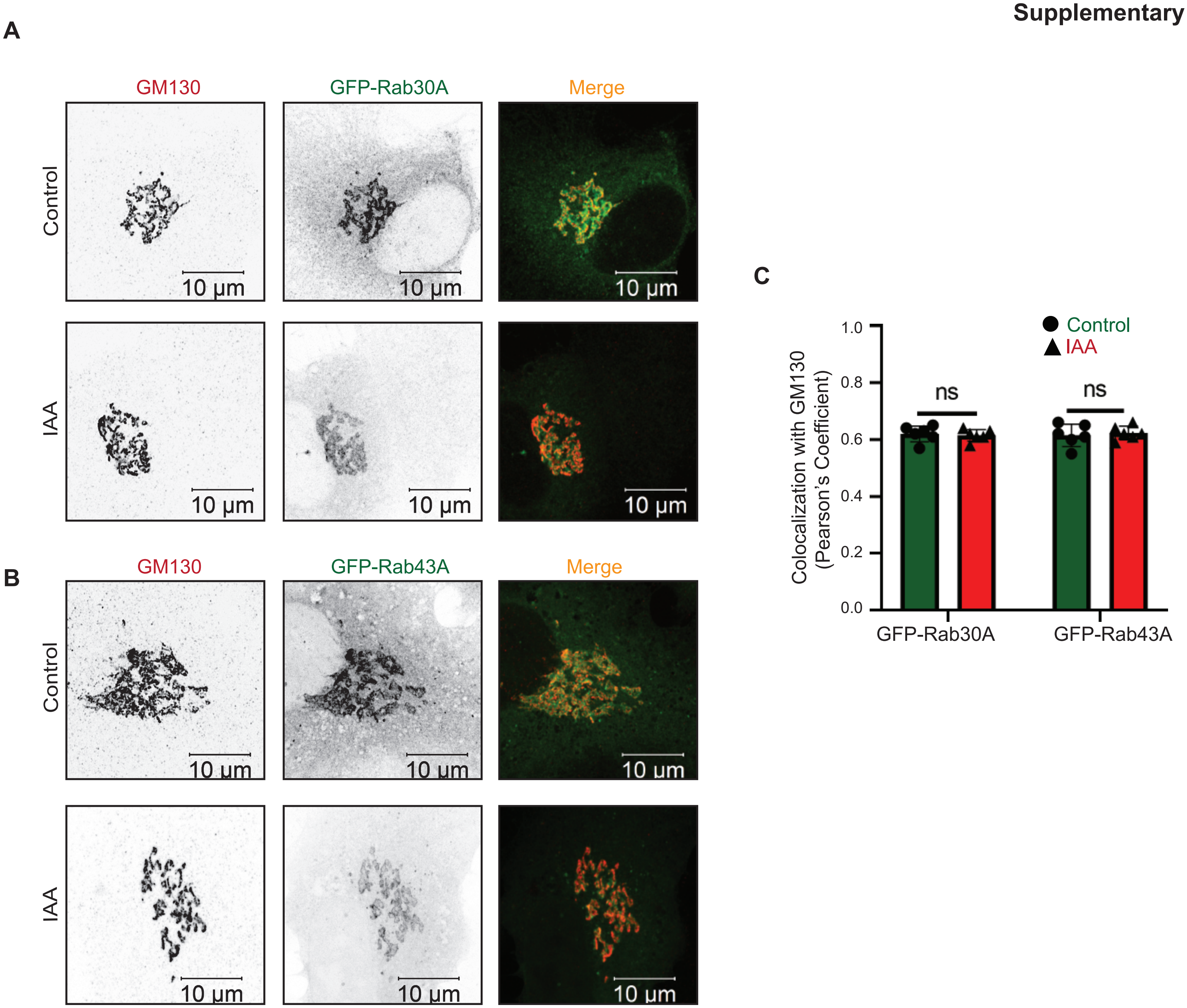

### Supplemental Figure 6

#### Supplementary 6

A

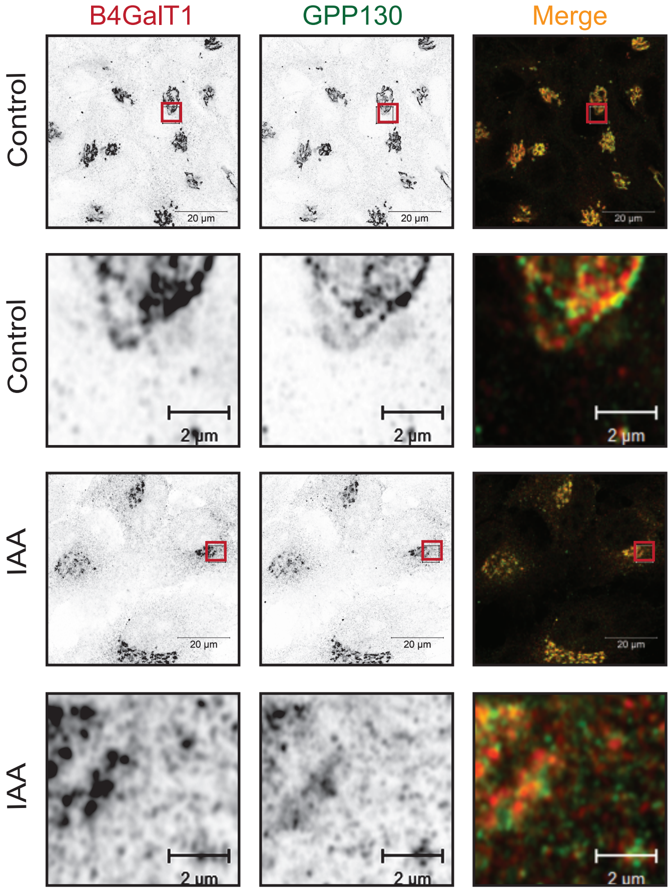


C

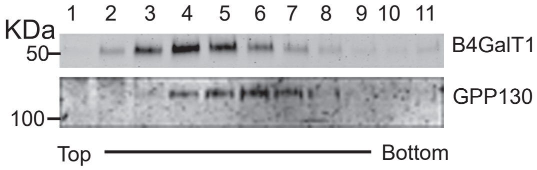


D

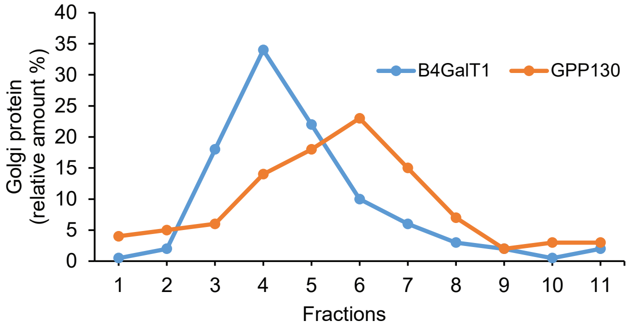


B

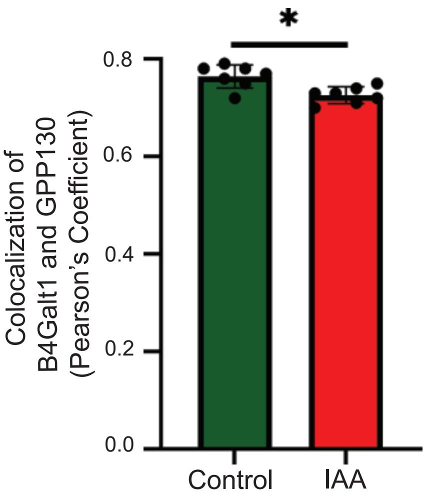

### Supplemental Figure 7

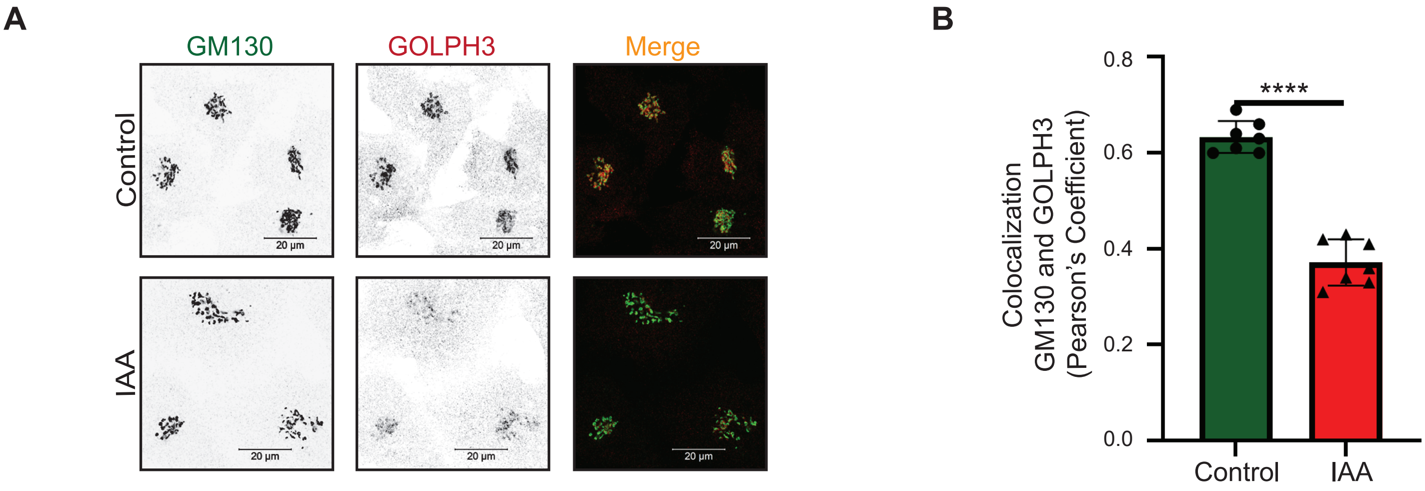
